# Super-enhancer driven expression of BAHCC1 promotes melanoma cell proliferation and genome stability

**DOI:** 10.1101/2023.01.18.524519

**Authors:** Pietro Berico, Maguelone Nogaret, Giovanni Gambi, Guillaume Davidson, Max Cigrang, Bujamin H Vokshi, Stephanie Le Gras, Gabrielle Mengus, Tao Ye, Mélanie Dalmasso, Emmanuel Compe, Corine Bertolotto, Irwin Davidson, Frédéric Coin

## Abstract

Super enhancers (SE) are stretches of active enhancers ensuring high expression levels of key genes associated with cell function and survival. The identification of cancer-specific SE-driven genes and their functional characterization may prove to be a powerful means for the development of innovative therapeutic strategies. By performing epigenomic profiling in patient-derived short-term melanoma cultures, we identify a SE promoting the specific expression of BAHCC1 in a broad panel of cutaneous and uveal melanoma cells. BAHCC1 is highly expressed in metastatic melanoma, correlates with decreased patient survival and is required for tumor growth. Integrative genomics analyses reveal that BAHCC1 is a transcriptional regulator controlling expression of a subset of E2F/KLF-dependent cell cycle and DNA repair genes. BAHCC1 associates with BRG1-containing remodeling complexes at the promoters of these genes. In agreement, BAHCC1 silencing leads to decreased cell proliferation and delay in DNA repair. Consequently, BAHCC1 deficiency cooperates with PARP inhibition to induce melanoma cell death. Our study identifies a novel SE-driven gene expressed in cutaneous and uveal melanoma and demonstrates how its inhibition can be exploited as a therapeutic target, alone or in combination with DNA damage-inducing agents.

## Introduction

Transcriptional deregulation represents a key mechanism for cancer initiation and progression (Bradner et al., 2017). A combination of somatic mutations and microenvironmental cues leads to the overexpression of epigenetic regulators promoting aberrant gene expression programs, ultimately resulting in cancer hallmarks (Hnisz et al., 2013). A key epigenetic mechanism promoting tumor-specific gene expression programs is the aberrant activation of Super-Enhancers (SEs), broad gene regulatory elements highly dependent on the activity of general co-activators compared to canonical enhancers (Pott and Lieb, 2015). Consequently, transcriptional inhibitors targeting co-activators such as BRD4 (Bromodomain-containing protein 4) and the cyclin-dependent kinase 7 (CDK7) of the basal transcription factor TFIIH are widely used to disrupt SE-associated genes in cancer cells (Filippakopoulos et al., 2010), (Kwiatkowski et al., 2014). However, the presence of SE-associated genes in normal cells, together with the poor pharmacokinetics and efficacy of transcriptional inhibitors in human clinical trials, underscore the need for alternative strategies (Postel-Vinay et al., 2016), (Ameratunga et al., 2020) such as the identification of cancer-specific SE-driven genes (Fontanals-Cirera et al., 2017).

Cutaneous melanoma remains the most lethal skin cancer whose incidence continues to increase over the past few decades (https://www.cancer.org/cancer/melanoma-skin-cancer/about/key-statistics.html). Despite the dramatic improvement in survival provided by therapies targeting mitogen-activated protein kinase (MAPK) coupled with the use of immune checkpoint inhibitors (Curti and Faries, 2021), many patients still develop resistance in part due to melanoma phenotypic plasticity, a dynamic and non-mutational mechanism of adaptation to micro-environmental changes and drug exposure (Marin-Bejar et al., 2021). Plasticity results in an important tumor heterogeneity involving multiple cell states with distinct transcriptional signatures and different proliferative, invasive and drug resistance phenotypes. Melanoma phenotype plasticity depends in part on two opposing gene expression programs, governed by the transcription factors MITF (microphthalmia-associated transcription factor) and SOX10 (SRY-box transcription factor 10) or AP-1 (Activating Protein 1) and TEAD (Transcriptional enhancer associate domain transcription factor), respectively, whose activities modulate melanoma cell state transition (Verfaillie et al., 2015), (Rambow et al., 2018), (Rambow et al., 2019), (Wouters et al., 2020). Other forms of melanoma include uveal melanoma, the most common primary intraocular tumor in adults that is intrinsically different from cutaneous melanoma. In uveal melanoma, the most frequent driver mutations are those involving the heterotrimeric G-protein subunits GNAQ or GNA11 (Onken et al., 2008), (Van Raamsdonk et al., 2010). Despite successful treatment of primary uveal melanoma, 50% of patients will develop metastasis that are highly refractory to existing treatments (Yang et al., 2018). Therefore, there is an urgent need to better understand the molecular mechanisms involved in these two cancers in order to develop efficient treatments.

Here, we characterized the SE landscapes of patient-derived short-term cutaneous melanoma cultures. Integrative epigenomic analyses revealed a melanoma-specific SE regulating the expression of the Bromo Adjacent Homology and Coiled Coil Domain-Containing 1 (BAHCC1) protein in a broad panel of cutaneous, but also in uveal melanoma cells. BAHCC1 drives cutaneous and uveal melanoma cell proliferation and is required for tumor growth of cutaneous melanoma xenografts *in vivo*. Loss-of-function and genomic profiling experiments show that BAHCC1 is a transcriptional regulator controlling the expression of a subset of E2F (E2 factor)/KLF (Krüppel-like factor)-dependent cell cycle and DNA repair genes in melanoma cells. BAHCC1 associates with BRG1 (BRM/SWI2-related gene 1)-containing remodeling complexes at the promoters of these genes. Consistent with the involvement of BAHCC1 in the regulation of DNA repair genes, including the crucial cell cycle kinase ATM (Ataxia Telengiectasia Mutated), BAHCC1 depletion delays DNA repair and cooperates with PARP (Poly ADP-ribose polymerase) inhibition to induce melanoma cell death. We thus identify a SE-dependent pan-melanoma expressed gene and demonstrate how its inhibition can be leveraged as a therapeutic target to impair melanoma cell proliferation, alone or in combination with DNA damage-inducing agents.

## Results

### SE17q25 regulates BAHCC1 expression in melanoma

To identify cutaneous melanoma (SKCM) specific SEs, we performed *in silico* ROSE analysis of H3K27ac ChIP-seq data from short-term melanoma cultures (MM) (Verfaillie et al., 2015) and two normal human melanocytes samples (NHEM1 and 2) (Fontanals-Cirera et al., 2017). The MM panel covered the two main phenotypes and most common driver mutations (Ext. Table 1). We further ranked SEs using DiffBind according to the enrichment in the binding of the lineage-specific transcription factors MITF and SOX10 that are recruited to long and short enhancers in melanocytic-like melanoma cells (Ext. Figure 1a) (Strub et al., 2011), (Laurette et al., 2015), (Berico et al., 2021), (Mauduit et al., 2021). Using these criteria, we identified a collection of potential SEs active in MM cells (Ext. Table 2). We further defined *bona fide* melanoma-specific SEs as active in at least 5 MM cell cultures. A SE at chr17q25.3 (hereafter referred to as SE17q25) met these criteria and was absent from NHEM (Figure 1a, left panel and Ext. Figure 1c). The SE17q25 region measured around 20kb and was recurrent not only in many of the MM cells, but also in melanoma cell lines such as SK-MEL-5 and 501mel (Figure 1a, left and right panels) (Ext. Table 2).

**Figure 1:**
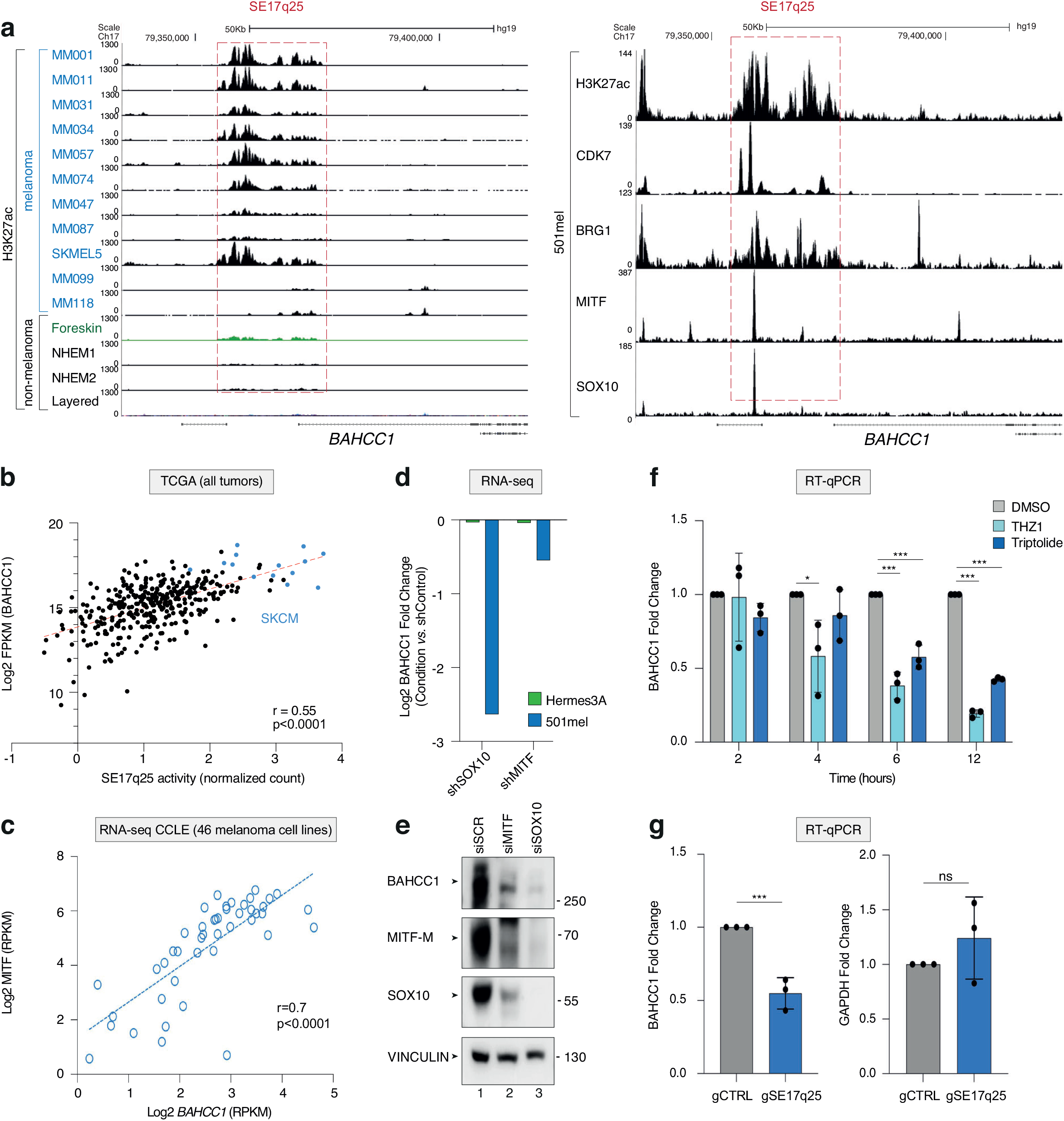
SE17q25 regulates BAHCC1 expression in melanoma. **a**. **Left panel**; Captures of the UCSC genome browser (GRCh38/hg19) showing the ChIP-seq profiles of H3K27ac in the genomic region of SE17q25 in several MM cell lines, normal melanocytes (NHEM = Normal Human Epidermal Melanocytes, Foreskin = Human Foreskin Melanocytes) and other tumor and normal cell lines (Layered = H3K27ac mark on 7 cell lines from ENCODE project). **Right panel**; Captures of the UCSC genome browser (GRCh38/hg19) showing the ChIP-seq profiles of H3K27ac, CDK7, BRG1, MITF and SOX10 at SE17q25 in mel501 cells. RefSeq annotated genes are displayed at the bottom. **b**. Spearman correlation between BAHCC1 RNA expression and SE17q25 activity, as defined by the ATAC-seq signal (normalized count), measured in different TCGA tumor samples (n=399). The SKCM samples are highlighted in light blue and have the higher expression of BAHCC1 and SE17q25 activity. “Spearman r” and p-value are shown on bottom right. **c**. Dot plot of BAHCC1 *vs*. MITF expression (RPKM) determined by RNA-seq from melanoma cell lines obtained from the CCLE data sets (n=49). The linear regression curve is shown in blue. “Spearman r” and p-value are shown on bottom right. **d**. BAHCC1 mRNA fold change (Condition *vs*. shControl) upon SOX10 or MITF KD in normal melanocytes (Hermes3A) or 501mel melanoma cells obtained from the GEO dataset GSE61967. **e.** 501mel melanoma cells were transfected with siSCR, siMITF and siSOX10 for 24 hours. Total extracts were resolved by SDS-PAGE and immuno-blotted against proteins as indicated. Molecular sizes are indicated. **f**. 501mel cells were treated with DMSO, THZ1 or Triptolide for 2, 4, 6 and 12 hours, as indicated and the relative amount of BAHCC1 mRNA was quantified by RT-qPCR. Bars represent mean values of three different experiments (Biological triplicates) (+/− SEM), Twoway ANOVA Šídák’s multiple comparisons test. **g.** RT-qPCR for GAPDH and BAHCC1 in 501mel cells co-transfected with CRISPR-dCas9^KRAB^ and guide RNAs (gRNA) against SE17q25 (gSE17q25) or a non-targeting genomic region (gCTRL). Bars represent mean values of three different experiments (Biological triplicates) (+/− SEM); paired t-test.

SE17q25 localized in close proximity to the promoter of the protein-coding gene BAHCC1 (Figure 1a). The Cancer Genome Atlas (TCGA) reports that high SE17q25 activity, measured by the level of ATAC-seq signal, correlated with high BAHCC1 expression in SKCM tumors (Figure 1b). Quantitative RT-PCR and Western blotting showed that BAHCC1 displayed higher RNA and protein levels in melanoma cells highly expressing MITF and SOX10, including melanocytic-like melanoma cells (501mel, MM117, IGR-37, SK-MEL-28, SK-MEL-5, MM011) and uveal melanoma (UVM) cells (OMM1, OMM1.3, OMM2.5) (Ext. Figures 1c and 1d). In agreement with these observations, analyses of the Cancer Cell Lines Encyclopedia (CCLE) revealed that BAHCC1 expression correlated with that of MITF in melanoma cell lines (n=49) (Figure 1c). Furthermore, depletion of SOX10 or MITF that decommissioned SE17q25 (Ext Figure 1e), strongly reduced BAHCC1 mRNA and protein expression in melanocytic-like melanoma cells (Figures 1d-e).

Compared to melanocytic-like melanoma cells, BAHCC1 was less expressed in dedifferentiated mesenchymal-like MITF^LOW^ melanoma cells (MM099, MM047, IGR-39 and MM029) (Ext. Figures 1c and 1d). Note, however, that BAHCC1 expression remained significantly higher in MITF^LOW^ melanoma cells compared to non-transformed Hermes3A melanocytes (which expresses MITF) or U-2 OS cells (non-melanoma cell line) (Ext. Figure 1d).

In addition to MITF and SOX10, the ATP-dependent chromatin remodeler BRG1 and the TFIIH kinase CDK7 that is known to associate with SEs (Chipumuro et al., 2014), also bound SE17q25 in 501mel cells (Figure 1a, right panel). Interestingly, treatment of 501mel cells for a short period of time with THZ1 (an inhibitor targeting CDK7) (Kwiatkowski et al., 2014) or Triptolide (an inhibitor of the XPB subunit of TFIIH) (Titov et al., 2011), (Noel et al., 2020) significantly diminished BAHCC1 expression (Figure 1f). In parallel, CRISPR interference (CRISPRi) of SE17q25 in 501mel using a dead Cas9 (dCas9) fused with the repressive Krüppel associated box (KRAB) domain (dCas9^KRAB^) and guide RNAs targeting SE17q25 (gSE17q25), diminished BAHCC1 expression compared to GAPDH control gene (Figure 1g). Altogether these data link SE17q25 activity to BAHCC1 expression in melanoma cells.

### BAHCC1 expression increases during melanoma progression

To explore BAHCC1 expression in tumors, we analyzed public transcriptional data and observed first that BAHCC1 was more strongly expressed in melanoma cell lines compared to other tumors cells (Figure 2a). In agreement, SKCM and UVM tumors from TCGA displayed higher levels of BAHCC1 compared to other tumors and normal tissues (Figure 2b). Furthermore, BAHCC1 overexpression was a marker of poor prognosis in SKCM or UVM patients (Figure 2c-d) and its expression was independent from the BRAF, NRAS or NF1 mutation status in SKCM patients (Figure 2e).

**Figure 2:**
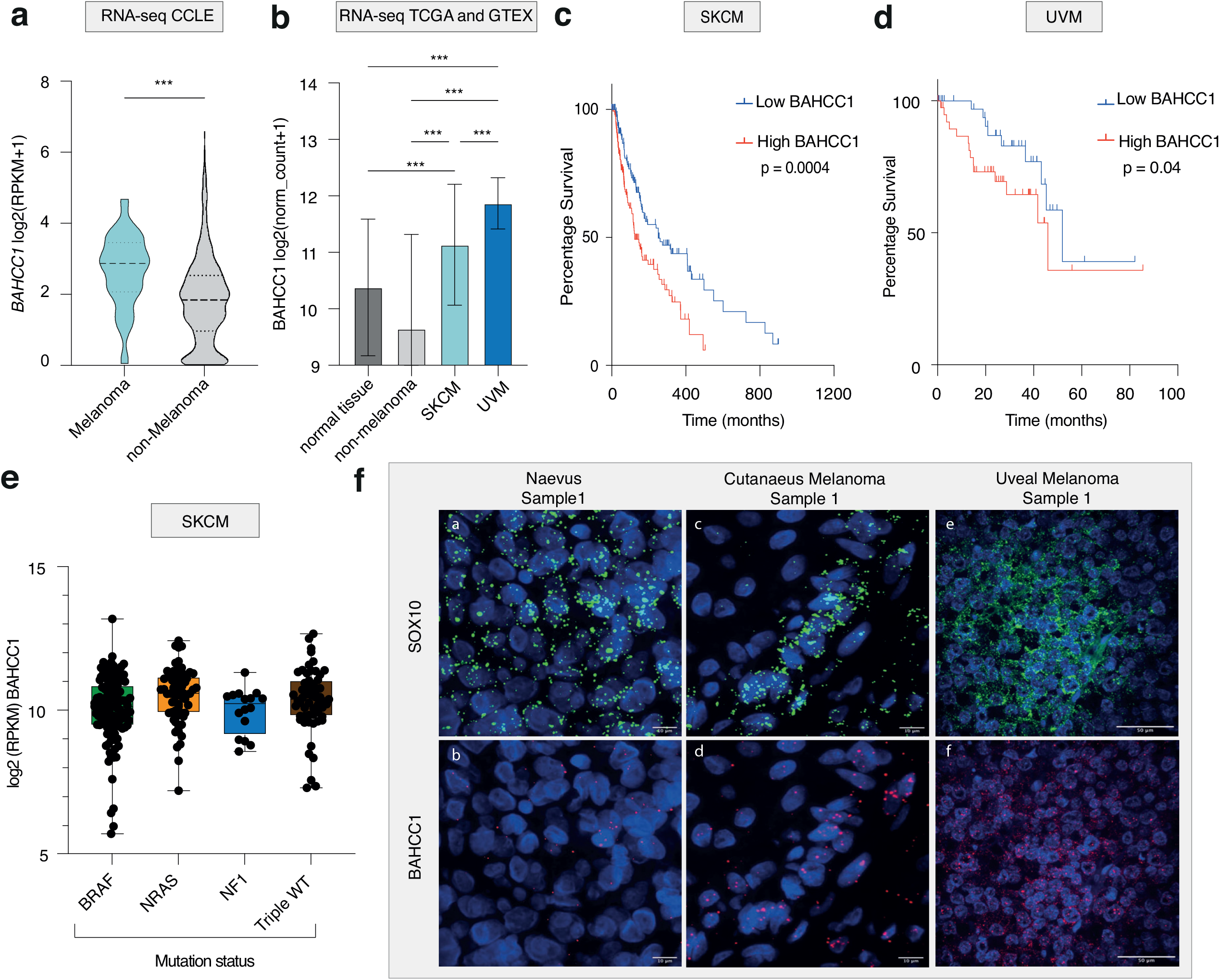
BAHCC1 is overexpressed in melanoma. **a**. Violin plot of BAHCC1 levels in melanoma *vs*. non-melanoma cell lines (RPKM) obtained from CCLE (n=1019); unpaired t-test. **b**. Normalized RNA levels of BAHCC1 in normal tissues (n=8156), non-melanoma tumors (n=10416), SKCM (n=469), UVM (n=79) obtained from TCGA and The Genotype-Tissue Expression (GTEX) datasets. **c-d**. Kaplan-Meier analysis of TCGA data from SKCM (n=302) **(c)** or UVM (n=80) **(d)** patients with high or low BAHCC1 expression (Lower and Upper percentile = 50); Log-rank Mantel-Cox test. **e**. Box and Whiskers plot representation of BAHCC1 expression (RPKM) in TCGA data from SKCM patients (n=469) according to the BRAF, NRAS and NF1 mutational status. Bars show the min and max values, box represent mean +/− SEM. **f**. RNA FISH against BAHCC1 and SOX10 mRNA in naevus, cutaneous and uveal melanomas tissue biopsies. Scale bars are indicated. Additional samples are shown in Ext. Figures 2e and f.

To trace the evolution of BAHCC1 expression along melanoma initiation and progression, different published bulk and single-cell transcriptomic datasets coming from normal cells, pre-malignant and malignant lesions were combined. Notably, we observed that BAHCC1 expression progressively increased going from normal skin to primary melanoma and metastatic lesions specifically in malignant cells (Ext. Figure 2a-d). Using RNA *in situ* hybridization on a cohort of patient samples including benign nevi, cutaneous and uveal melanoma, we confirmed the increased expression of BAHCC1 mRNA in SOX10-positive malignant cells (Fig 2f and Ext. Figure 2e-f). Overall, these results indicate that BAHCC1 expression progressively increases during melanoma progression from primary tumors to metastasis.

### BAHCC1 is required for melanoma cell proliferation and tumor growth

To investigate the functional consequences of BAHCC1 knock down (BAHCC1 KD), we transfected melanoma cells with two independent custom antisense oligonucleotides (ASO, locked nucleic acid GapmeRs) targeting the pre-mRNA. GapmeRs #1 and #2 efficiently silenced BAHCC1 expression in 501mel cells (Ext. Figure 3a). We observed that BAHCC1 KD reduced proliferation of cutaneous melanocytic-like (501mel and MM117) and mesenchymal-like (MM047 and MM029) melanoma cells but also of uveal melanoma cell lines (OMM1.3 and OMM2.5) (Figure 3a, Ext. Figure 3b). In contrast, BAHCC1 KD did not affect proliferation of U-2 OS cells excluding an off-target effect (Ext. Figure 3b). Given the anti-proliferative role of MAPKi inhibitors (MAPKi) used in the clinic, we wondered whether BAHCC1 KD could increase their effect. Indeed, BAHCC1 KD further reduced the proliferation of a melanocytic cell line upon treatment with Trametinib or Vemurafenib (Figure 3b). We also evaluated the effect of BAHCC1 KD on melanoma cell invasive capacity and observed that BAHCC1 KD significantly impaired migration of 501mel, MM099 and MM029 melanoma cells (Figure 3c). All together, these results highlight the importance of BAHCC1 for melanoma cells proliferation and invasion *in vitro*.

**Figure 3:**
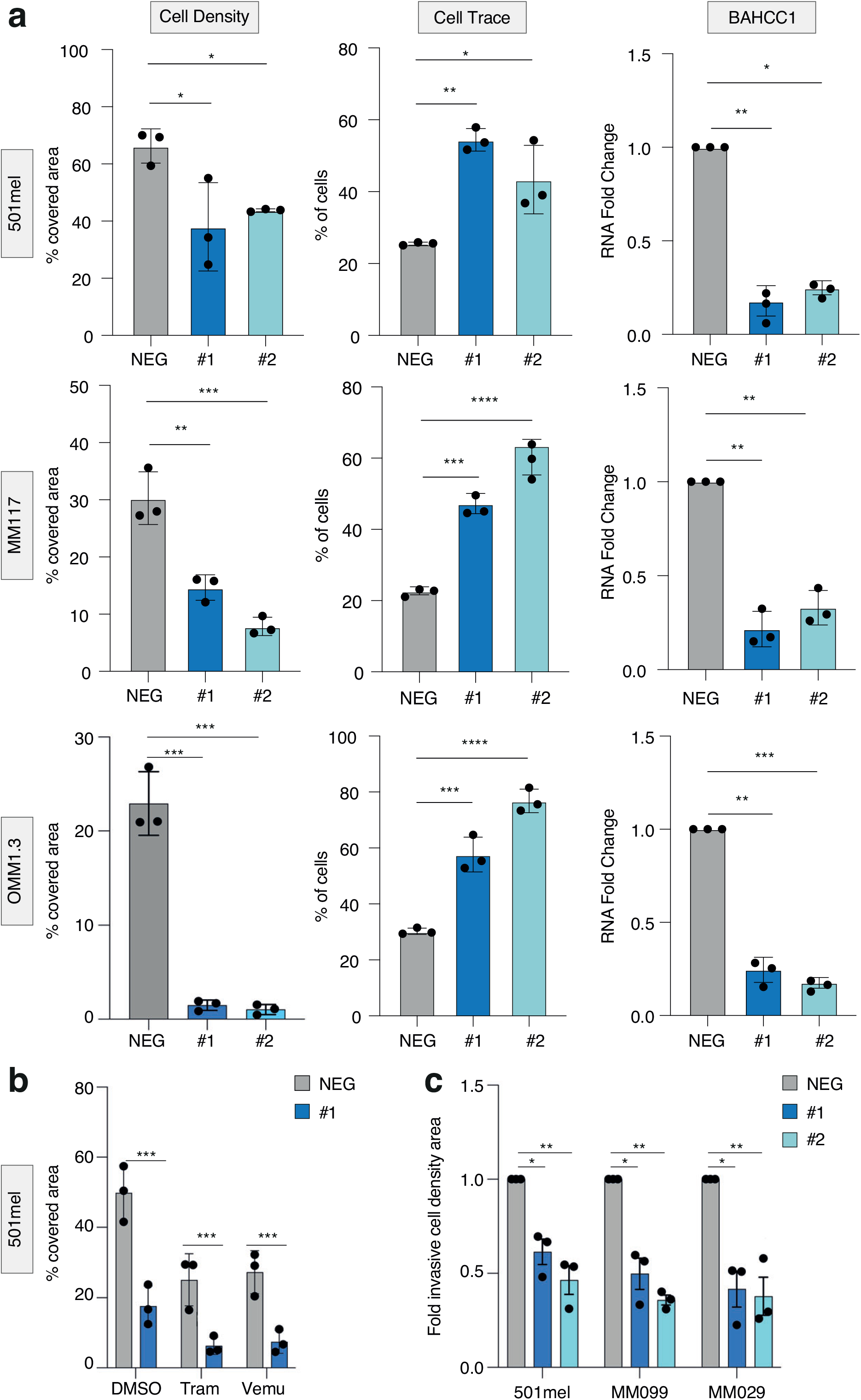
BAHCC1 depletion impairs melanoma cell proliferation. **a**. **Left panel**; Quantification of crystal violet staining of cells transfected with GapmeR^NEG^ (NEG), GapmeR^#1^ (#1) and GapmeR^#2^ (#2) in SKCM cells (501mel and MM117) and UVM (OMM1.3) cells. **Middle panel**; CellTrace staining was measured by FACS in the cells used in the left panel and results are represented as % of slow proliferative cells considering an arbitrary threshold between 20-30% in the GapmeR^NEG^ (NEG) control. **Right panel**; Relative BAHCC1 expression upon transfection with GapmeR^NEG^ (NEG), GapmeR^#1^ (#1) and GapmeR^#2^ (#2) were measured by RT-qPCR in the cells used in the left panel. Bars represent mean values of three different experiments (Biological triplicates) (+/− SEM). Ordinary oneway ANOVA using Dunnett’s multiple comparisons test. **b**. Cell coverage quantification of crystal violet staining of 501mel cells transfected with GapmeR^NEG^ (NEG) or GapmeR^#1^ (#1) upon treatment with DMSO, Trametinib (10nM) or Vemurafenib (5uM). Bars represent mean values of three different experiments (Biological triplicates) (+/− SEM); Ordinary one-way ANOVA using Dunnett’s multiple comparisons test. **c**. Cell coverage quantification of the Boyden-Chamber transwell after crystal violet staining of 501mel, MM099 and MM029 cells transfected with GapmeR^NEG^ (NEG), GapmeR^#1^ (#1) and GapmeR^#2^ (#2). Bars represent mean values of three different experiments (Biological triplicates) (+/− SEM); Two-way ANOVA using Tukey’s multiple comparisons test.

To test the effects of BAHCC1 silencing *in vivo*, we injected the MITF^HIGH^ IGR37 melanoma cell line expressing the Firefly Luciferase together with a doxycycline-inducible anti-BAHCC1 or non-targeting shRNA (IGR37^shBAHCC1^ and IGR37^shCTRL^, respectively) in nude mice (Ext Figure 4a-c). ShRNA-mediated KD of BAHCC1 significantly suppressed CDX tumour growth compared to the non-targeting shRNA (Figure 4a). These results were consistent with a reduced luminescence from the IGR37^shBAHCC1^-derived tumors, compared to the control IGR37^shCTRL^ (Figure 4b and Ext Figure 4d). Immunoblots performed on tumor samples confirmed the strongly diminished BAHCC1 protein level in tumors derived from IGR37^shBAHCC1^ cells (Figures 4c-d). These data point to the critical role of BAHCC1 in melanoma cell proliferation and tumor growth *in vivo*.

**Figure 4:**
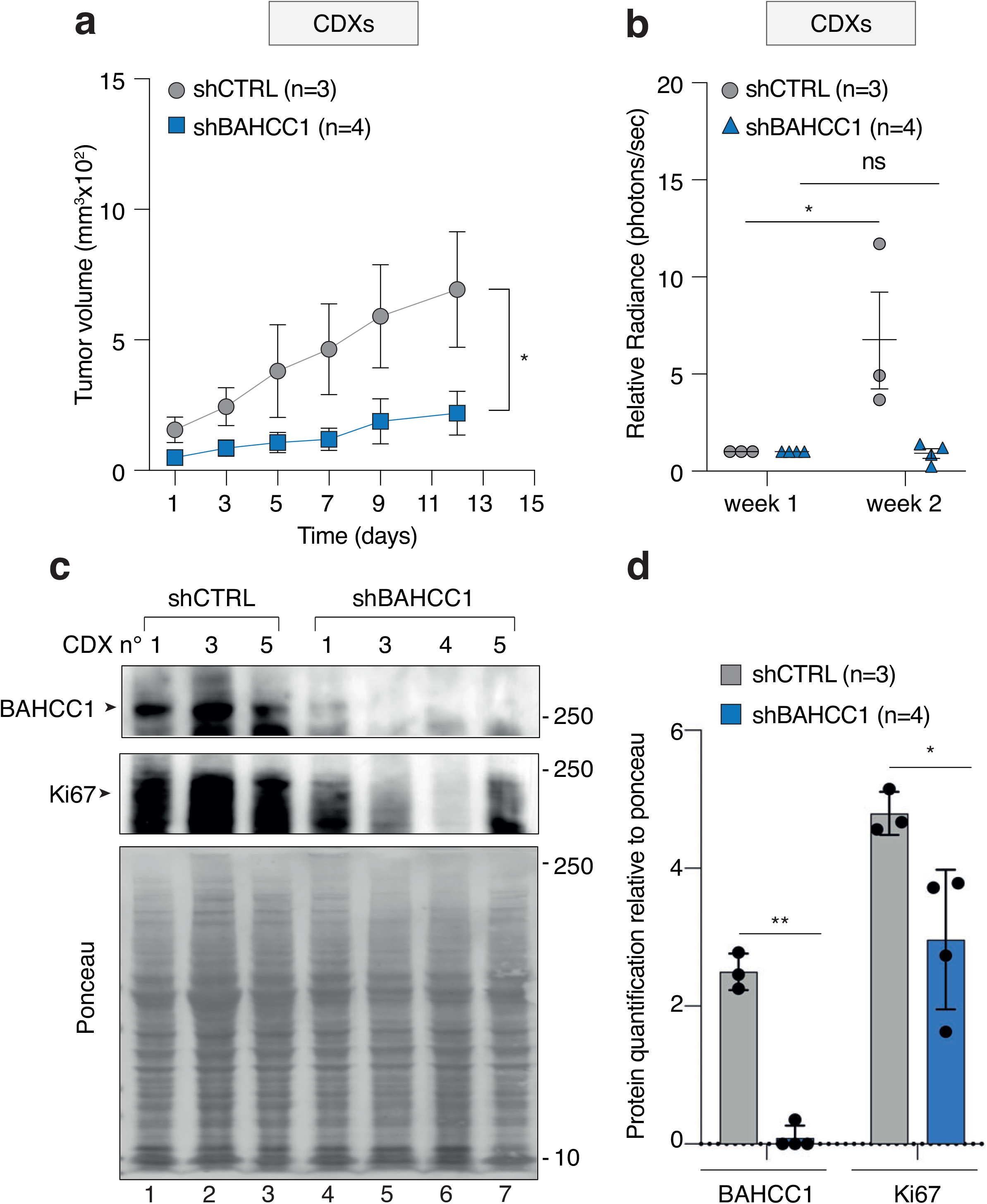
BAHCC1 depletion impairs melanoma tumor growth. **a**. Kinetic of tumor growth obtained following xenografting of IGR37 melanoma cells stably expressing inducible shCTRL or shBAHCC1 in nude mice (+/− SEM); Holm-Sídák’s multiple comparisons test. **b.** Relative luciferase activity expressed as radiance of the tumors measured after one week of growth. Each point corresponds to one mouse; One-way ANOVA paired test. **c**. Total extracts of tumors obtained following xenografting of IGR37 melanoma cells stably expressing inducible shCTRL or shBAHCC1 in nude mice were resolved by SDS-PAGE and immunoblotted against proteins as indicated. Ki67 is a marker of cell proliferation. Molecular sizes are indicated. **d**. Immunoblot signals of panel (c) were quantified and average BAHCC1 and Ki67 protein amount, relative to ponceau, was reported on the graph (+/−SEM); Šídák’s multiple comparisons test.

### BAHCC1 regulates E2F/KLF target genes involved in DNA repair and cell cycle

To better understand BAHCC1 function in melanoma cell proliferation, we first checked its localization by immunofluorescence in 501mel cells and observed its nuclear localization (Ext Figure 5a). Analysis of subcellular protein fractions by immunoblot confirmed BAHCC1 accumulation in the nuclear soluble and insoluble fractions (Ext Figures 5b). We then investigated the potential role of BAHCC1 to regulate melanoma gene expression first by transcriptome profiling of 501mel and MM047 cells before and after BAHCC1 KD. RNA-seq revealed that several hundred genes were deregulated in the absence of BAHCC1 (absolute log2 fold change >0.5 and adjusted P-value<0.05) (Figure 5a) with a significant overlap of 200 genes between the 501mel and MM047 cells (Figure 5b and Ext. Table 3). Gene Ontology (GO) analyses of the 200 co-downregulated and 82 co-upregulated genes revealed their enrichment in cell cycle (MCM7, TOP2A, CNTRL), DNA repair (RAD51B, FANCG, ATM), regulation of MAPK-pathway (MAP2K1, CDKN1B, CDKN2C) as well as mitochondria biogenesis (MTFR1, IMMP2L, DNA2) and cell migration (MMP2, COL18A1, BMP1) (Ext Figures 5c), consistent with the anti-proliferative and anti-invasive effects of BAHCC1 KD observed above.

**Figure 5:**
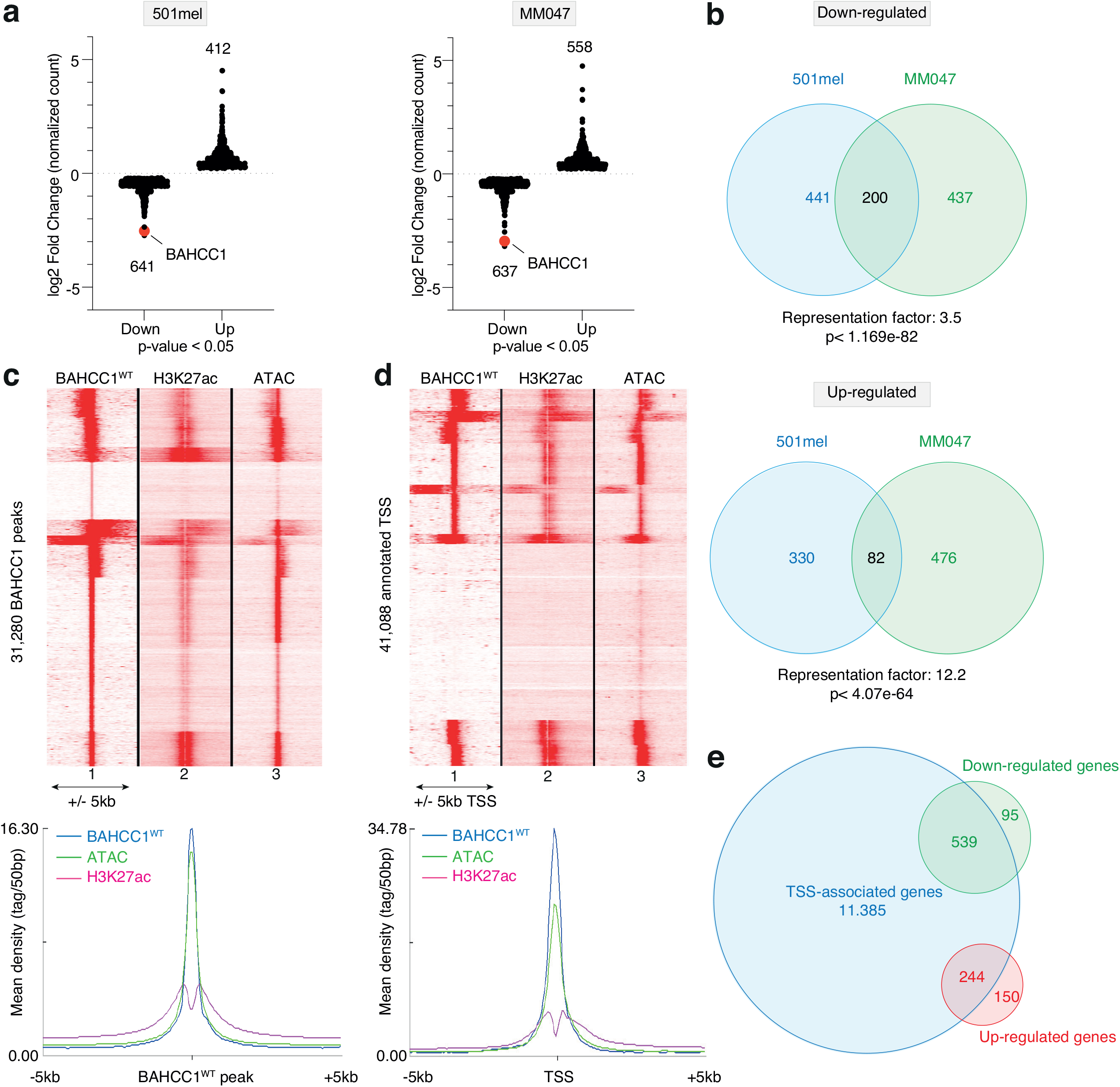
BAHCC1 is a transcriptional regulator. **a**. Scatter plot of the significantly (p-value<0.05) deregulated genes (normalized count) upon BAHCC1 KD in 501mel and MM047. Red dots highlight BAHCC1 which is one of the top downregulated genes. **b**. Venn diagram between significantly down-regulated (top) and up-regulated (bottom) genes identified by RNA-seq in 501mel and MM047 upon BAHCC1 KD. Representation factor and p-values were calculated using hypergeometric test. **c**. **Upper panel**; Read density clustering obtained with seqMINER for the 31,280 BAHCC1 occupied sites relative to BAHCC1, H3K27ac and ATAC-seq signals in 501mel cells in a genomic window of 10 kb around the peaks. **Lower panel**; Merge meta-profiles distribution of BAHCC1, H3K27ac and ATAC enrichment relative to the 31,280 BAHCC1 peaks. **d**. **Upper panel**; Read density clustering obtained with seqMINER for BAHCC1, H3K27ac and ATAC-seq signals relative to the 41,088 annotated TSS. **Lower panel**; Merge meta-profiles distribution of BAHCC1, H3K27ac and ATAC in a +/− 5 kb window around the TSS. **e**. Venn diagram between genes showing a TSS-associated BAHCC1 protein and the significantly down- or up-regulated genes following BAHCC1 KD, as determined by RNA-seq.

To evaluate if BAHCC1 directly regulated gene expression by chromatin binding, we profiled endogenous BAHCC1 (BAHCC1^WT^) genome-wide occupancy using CUT&Tag followed by deep sequencing in 501mel cells. We identified 31,280 peaks for endogenous BAHCC1^WT^, localizing mostly in proximal gene promoters (37.62%) and distal intergenic (24.28%) regions (Ext Figures 5d). Read density clustering of BAHCC1^WT^, H3K27ac ChIP-seq and ATAC-seq in the 31,280 BAHCC1-occupied regions demonstrated that BAHCC1^WT^ mostly binds to active chromatin at the nucleosome depleted regions between two H3K27ac-marked nucleosomes (Figure 5c). Moreover, a large fraction of the 41,088 annotated transcription start-sites (TSS) showed a TSS-centered BAHCC1 and ATAC peaks flanked by H3K27ac chromatin marks (Figure 5d). Consistent with these data, analysis of the 1,000 best scoring BAHCC1-occupied sites with RSAT to identify transcription factor DNA binding motifs and transcription factors that may cooperate with or recruit BAHCC1 to regulate gene expression revealed strong enrichment of the NFY and SP family factors known to be enriched at proximal promoters (Ext. Table 4). Integrating BAHCC1 RNA-seq and CUT&Tag data, we observed that amongst the 11,385 genes associated with a TSS-centered BAHCC1 peaks, 783 genes were deregulated upon BAHCC1 KD in 501mel cells, of which 539 (70%) were down-regulated and 244 (30%) were up-regulated (Figure 5e). RSAT analyses of the DNA motifs under the BAHCC1 bound sites at the down-regulated promoters showed enrichment of NFY as above but also for motifs for KLF-family factors. Analyses at the promoters of up-regulated genes also showed KLF-family factors, but also SOX10 (Ext. Table 4).

Further evidence for the role of BAHCC1 in cell cycle gene regulation came from mining of single cell RNA-sequencing (scRNA-seq) from melanoma patient derived xenografts (PDXs) (Rambow et al., 2018), (Berico et al., 2021) where BAHCC1 was preferentially expressed in the “mitotic” melanoma cell subpopulation (Ext Figures 5e) also significantly enriched in E2F and SP/KLF regulons (Ext Figures 5f). Similarly, scRNA-seq from uveal and cutaneous tumors (Tirosh et al., 2016), (Pandiani et al., 2021) were separated into “Slow” or “Fast” cycling cells according to the Tirosh cell cycle signature (95 genes) (Ext. Table 5) (Tirosh et al., 2016). We observed that BAHCC1 was significantly enriched in fast cycling cells (Ext Figures 5g), confirming its potential role in the control of the melanoma cell cycle. Similar observations were made using TCGA datasets from UVM or SKCM patients, in which BAHCC1 expression significantly correlated with the Tirosh cell cycle signature (Ext Figures 5h). Together, these results demonstrate that BAHCC1 regulates the expression of a set of E2F/KLF-dependent genes involved in cell cycle and DNA repair.

### BAHCC1 interacts with BRG1-containing complexes

Protein Blast and AlphaFold tools revealed the presence of a Coiled Coil (CC) region in the central part of the protein together with two well conserved TUDOR and Bromo-Adjacent Homology (BAH) domains at the C-terminal (Figure 6a). CCs are involved in protein-protein interactions, while BAH and TUDOR domains are found in a wide range of chromatin binding proteins where they are readers of histone modifications and act as chromatin co-activators/repressors (Ciani et al., 2010), (Musselman et al., 2012). We profiled genome occupancy of a FLAG-tagged truncated form of BAHCC1 missing the N-terminal CC domain and bearing only the short C-terminal region containing the TUDOR and BAH domains (BAHCC1^TUDOR-BAH^) (Figure 6a). Genome-wide BAHCC1^TUDOR-BAH^ occupancy pattern highly overlapped with endogenous BAHCC1^WT^ as shown by seqMINER read density heatmap (Figure 6b), further validating the CUT&Tag data obtained above with the full-length endogenous protein. However, a limited cluster of sites was selectively occupied by endogenous BAHCC1^WT^, but not the truncated form (Figure 6b and Ext. Table 6). Importantly, this BAHCC1 N-terminal region-dependent cluster included 222 of the 539 genes down-regulated after BAHCC1 KD, including DNA repair or cell cycle genes such as ATM or CDKN1A (Figure 6c, Ext Figures 6a and Ext. Table 6). In agreement with this, RSAT analyses showed that N-terminal region-dependent sites were preferentially enriched in KLF-family DNA binding motifs similar to the down-regulated promoters (Ext. Table 4).

**Figure 6:**
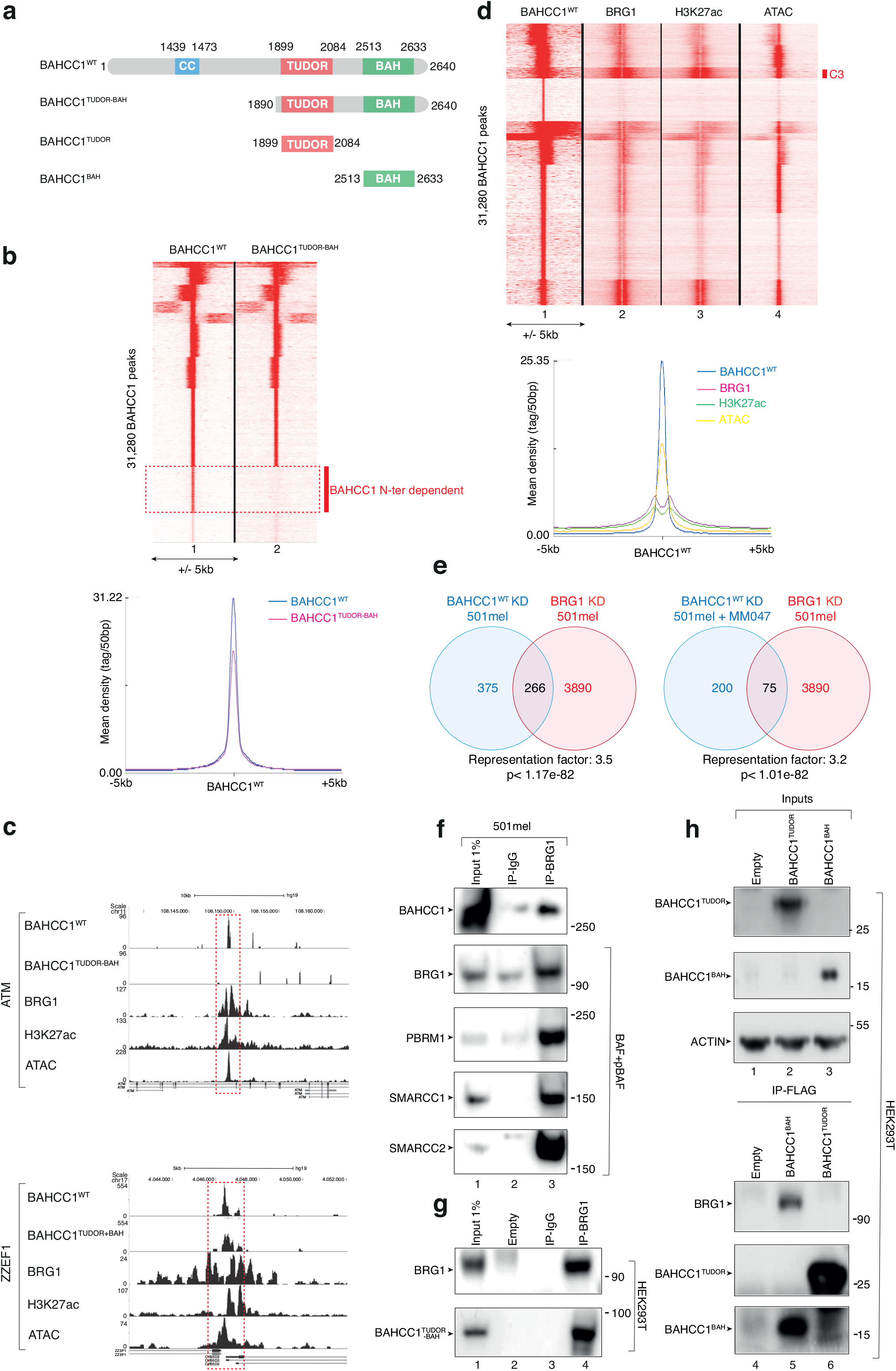
BAHCC1 interacts with BRG1-containing chromatin remodeler complexes. **a**. Domain architecture of BAHCC1. CC; Coiled Coil. The deletion mutants used below are shown. **b**. **Upper panel**; Read density clustering using SeqMINER showing the colocalization between BAHCC1^WT^ and BAHCC1 ^TUDOR-BAH^ in the 31,280 BAHCC1^WT^ occupied sites. Red box indicates the cluster of BAHCC1 N-terminal region dependent peaks. **Lower panel**; Metaprofiles of BAHCC1^WT^ and BAHCC1 ^TUDOR-BAH^ signals around the 31,280 BAHCC1 peaks. **c.** Captures of the UCSC genome browser (GRCh38/hg19) showing the ChIP-seq profiles for BAHCC1^WT^, BAHCC1 ^TUDOR-BAH^, BRG1, H3K27ac and ATAC-seq in 501mel corresponding to the promoter regions of ATM and ZZEF1 genes. ZZEF1 is used as a control gene recruiting both BAHCC1^WT^ and BAHCC1^TUDOR-BAH^. RefSeq annotated genes are displayed at the bottom. **d**. **Left panel**; Read density clustering using SeqMINER showing the colocalization between BAHCC1^WT^, BRG1, H3K27ac and ATAC signal in the 31,280 BAHCC1-occupied sites. Cluster 3 (C3) is highlighted. **Right panel**; Meta-profile of BAHCC1, BRG1, H3K27ac and ATAC signals around the 31,280 BAHCC1 peaks. **e**. **Left panel:** Venn diagram showing the overlap between the down regulated genes in 501mel following either BRG1 or BAHCC1 KD. **Right panel** Venn diagram showing the overlap between the down-regulated genes in mel501 after BRG1 KD and the common down-regulated genes in 501mel and MM047 after BAHCC1 KD. Representation factor and p-values were calculated using hypergeometric test. **f**. BRG1 was immunoprecipitated (IP-BRG1) from nuclear extracts of 501mel cells. Following SDS-PAGE, proteins were immunoblotted as indicated, including subunits of the pBAF/BAF complexes. IP-IgG was performed as a negative control. Molecular sizes are indicated. **g**. HEK293T cells were transfected to express HA-BAHCC1^TUDOR-BAH^ protein and BRG1 was immunoprecipitated (IP-BRG1). Following SDS-PAGE, proteins were immunoblotted as indicated. IP-IgG was performed as a negative control. Molecular sizes are indicated. **h**. HEK293T cells were transfected to express FLAG-BAHCC1^BAH^ and FLAG-BAHCC1^TUDOR^ domains and FLAG-IP was performed. Following SDS-PAGE, proteins were immunoblotted as indicated. Molecular sizes are indicated.

We previously demonstrated that the BRG1 subunit of the PBAF chromatin remodeling complex occupied H3K27ac marked nucleosomes in 501mel melanoma cells (Laurette et al., 2015). Comparison of genome wide occupancy of both BRG1 and BAHCC1 showed that most BAHCC1 peaks were flanked by nucleosomes bound by BRG1 and marked by H3K27ac (Figure 6d). SeqMINER read density clustering allowed the identification of several BAHCC1-BRG1 clusters amongst which cluster 3 (C3) was associated with DNA repair and cell cycle genes including ATM that was strongly enriched in BAHCC1, BRG1 and H3K27ac at its promoter (Figure 6c). Moreover, a significant fraction of genes down-regulated upon BAHCC1 KD in 501mel were also down-regulated by BRG1 silencing (Figure 6e, left panel) including 75 of the 200 genes down-regulated in both 501mel and MM047 (Figure 6e, right panel). These genes were found to be mostly involved in mitosis and DNA repair, including ATM (Ext Figures 6b and Ext. Table 3).

We next aimed to investigate whether a physical interaction could be observed between BAHCC1 and BRG1-containing complexes. BRG1 co-immunoprecipitated with endogenous BAHCC1, together with the BAF and pBAF subunits PBRM1, SMARCC1 and SMARCC2 in 501mel cells (Figure 6f). In parallel, pull down of BRG1 co-precipitated the FLAG-tagged BAHCC1^TUDOR-BAH^ deletion mutant overexpressed in HEK293T cells (Figure 6g). We further observed that BRG1 co-precipitated specifically with the BAH domain of BAHCC1 but not with the TUDOR domain (Figure 6h). The BAHCC1^BAH^-BRG1 co-precipitation was confirmed in 501mel melanoma cells stably expressing HA-tagged BAHCC1^BAH^ domain (Ext Figures 6c). Altogether, these data highlight a direct physical interaction between BAHCC1 and BRG1-containing BAF and PBAF complexes that regulate a common subset of genes.

### BAHCC1 KD cooperates with PARP inhibition to induce melanoma cell death

The above results suggested a role of BAHCC1 in genome stability through its regulation of DNA repair proteins, including the master DNA damage repair sensor ATM. In agreement, RT-qPCR and immunoblotting showed that BAHCC1 KD decreased expression of ATM at the mRNA and protein levels (Figures 7a and b). Furthermore, BAHCC1 expression positively correlated with genome alteration frequencies in human melanoma tumors (Figure 7c). We then tested the effect of BAHCC1 KD on the ability of 501mel cells to repair DNA following treatment with Neocarzinostatin (NCS) that induces DNA double strand breaks (Figure 7d). Strikingly, BAHCC1 KD led to higher numbers of γH2AX foci during the repair time course, indicating a delay in the repair of DNA double strand breaks in the absence of BAHCC1 (Figure 7e and Ext Figure 7).

**Figure 7:**
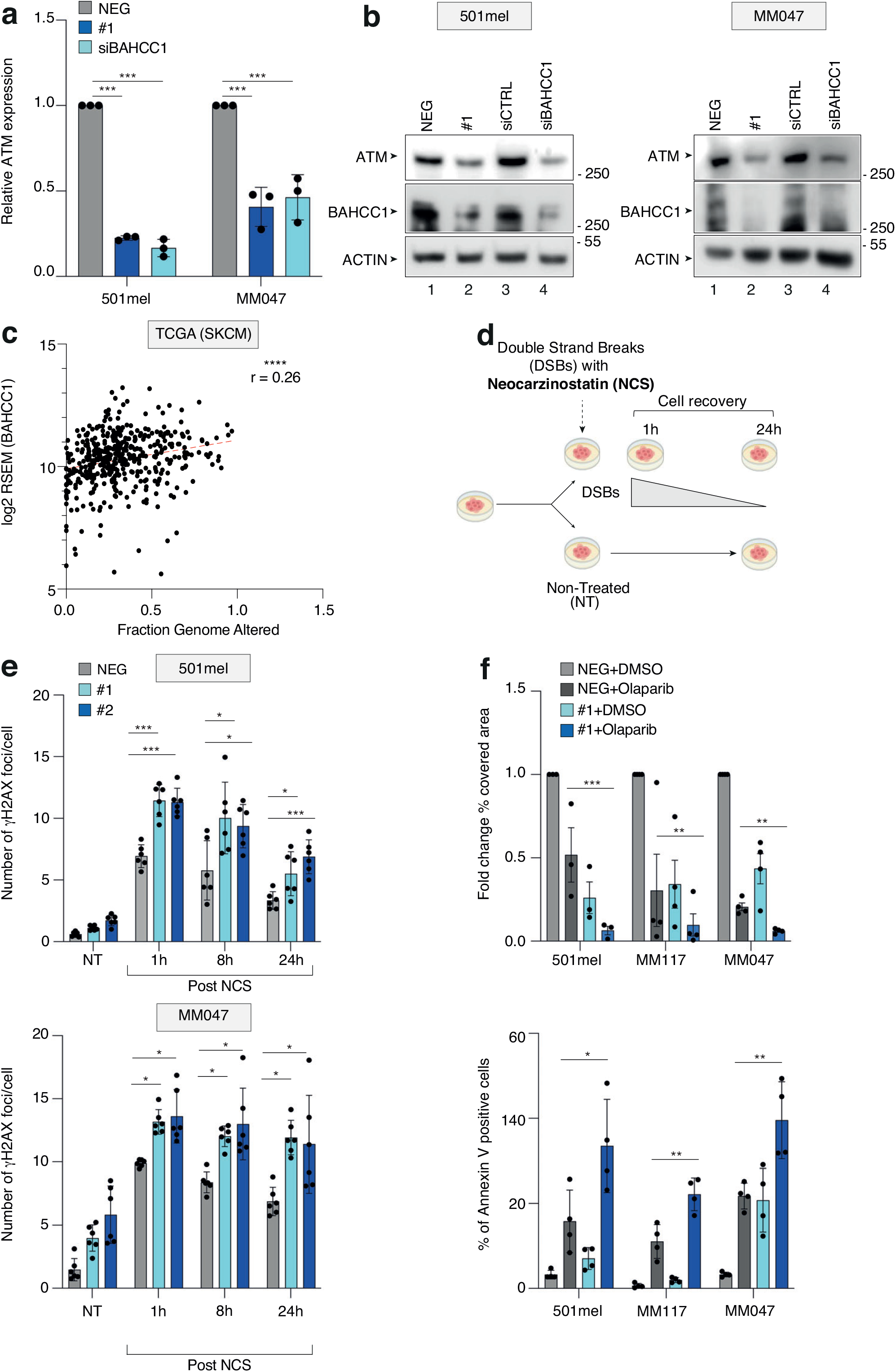
BAHCC1 depletion cooperates with PARPi to induce cell death. **a**. Relative ATM expression upon transfection with GapmeR^NEG^ (NEG), GapmeR^#1^ (#1) and shBAHCC1 was measured by RT-qPCR in 501mel or MM047. Bars represent mean values of three different experiments (Biological triplicates) (+/− SEM); two-way ANOVA using Šídák’s multiple comparisons test. **b**. 501mel or MM047 were transfected with GapmeR^NEG^ (NEG), GapmeR^#1^ (#1), shCTRL or shBAHCC1. Whole cell extracts were resolved by SDS-PAGE and proteins were immunoblotted as indicated. **c.** Spearman correlation between BAHCC1 expression and the fraction of genomic alteration (FGA) in TCGA melanoma samples (n=443) where FGA is considered as the percentage of copy number alterations found in the tumor compared to the normal karyotype. The linear regression curve is shown in red. **d**. Schematic representation of the *in vitro* experiments using Neocarzinostatin (NCS). **e**. Immunofluorescence quantification of the number of γH2AX foci per cell in 501mel and MM047 transfected either with GapmeR^NEG^ (NEG), GapmeR^#1^ (#1) or GapmeR^#2^ (#2) and treated or not (NT) with NCS (1, 8 and 24 hours of recovery). Bars represent the means obtained from six biological replicates (+/−SEM) (501mel n1=45338, n2=46604, n3=47365, n4=39185, n5=61975, n6=85783; MM047 n1=26535, n2=26284, n3=28105, n4=8559, n5=13342, n6=18296); Two-way ANOVA test. **f**. **Left panel**; Crystal violet quantification expressed as fold change relative to GapmeR (NEG)-transfected cells treated with DMSO. Bars represent mean values of three different experiments (Biological triplicates) (+/− SEM); Two-way ANOVA using Dunnett’s multiple comparisons test. **Right panel**; Percentage of Annexin V positive cell in 501mel, MM117 and MM047 transfected with GapmeR^NEG^ (NEG) and GapmeR^#1^ (#1) and treated for 96 hours with DMSO or 10uM of Olaparib. Bars represent mean values of three different experiments (Biological triplicates) (+/− SEM); Two-way ANOVA using Dunnett’s multiple comparisons test.

Poly (ADP-ribose) polymerase 1 (PARP1) is the main PARP-family protein involved in DNA damage response where it acts as a DNA damage sensor. PARP1 inhibition (PARPi) has been shown to increase DNA replication fork destabilization leading to accumulation of DNA breaks (Caron et al., 2019). Since the PARP-pathway acts independently from ATM, it has been postulated that ATM-deficient cancer cells become addicted to PARP-dependent DNA repair, making them highly sensitive to PARPi (Pilié et al., 2019). Therefore, we queried whether BAHCC1 KD might potentiate the effect of PARPi in melanoma cells. Strikingly, cotreatment with the PARPi Olaparib and BAHCC1 KD showed a significant cooperative effect on viability of 501mel (MITF^HIGH^, BRAF^V600E^), MM117 (MITF^HIGH^, Triple-wt) and MM047 (MITF^LOW^, NRAS^Q61R^) melanoma cells due to increased apoptosis (Figure 7f). Overall, these findings demonstrate the important role of BAHCC1 in the expression of DNA repair genes such as ATM, the kinetics of DNA-repair and are in line with the idea that melanoma cells may be sensitized to PARPi by BAHCC1 depletion.

## Discussion

Deregulation of gene expression in cancer cells is well established and cannot be explained solely by genomic alterations such as mutations or copy number variations. Cancer cells undergo significant changes in their transcriptional program through extensive rewiring that includes the acquisition of alternative gene regulatory elements such as SEs (Lee and Young, 2013), (Lovén et al., 2013), (Bradner et al., 2017). Here, using several H3K27ac ChIP-seq data sets from short-term patient-derived cutaneous melanoma cultures, we identified melanoma-specific SEs and their associated genes that could also be exploited in therapy. We further integrated the H3K27ac profile with the binding profiles of master regulators in melanoma cells such as SOX10 and MITF. These transcription factors occupy long and short enhancers found in cutaneous melanocytic-like melanoma cells with SOX10 being required to achieve high levels of activity (Mauduit et al., 2021). Our analysis converged on the SE17q25 element that was activated in most of the melanocytic-like melanoma cells but also in SKCM and UVM biopsies regardless of driver mutation status (BRAF, NRAS, Triple-WT in cutaneous and GNAQ or GNA11 in uveal melanoma). In cutaneous melanoma cells, the SE17q25 element was not only highly occupied by MITF and SOX10, but also by the TFIIH kinase CDK7 and BRG1, all of which are known to occupy relevant SEs (Kwiatkowski et al., 2014), (Barutcu et al., 2016), (Eliades et al., 2018). Depletion of MITF or SOX10 as well as TFIIH inhibition decommissioned SE17q25 and reduced the expression of BAHCC1, a gene located in close vicinity to SE17q25. In addition, selective CRISPR-mediated silencing of SE17q25 significantly affected BAHCC1 expression strongly supporting the idea that SE17q25 regulated BAHCC1 expression.

In agreement with its dependency on MITF and SOX10 observed *in vitro*, high BAHCC1 expression correlated with high MITF and SOX10 expression in melanoma biopsies and with poor prognosis in both SKCM and UVM patients (Gao et al., 2020). Consistent with these findings, an extensive analysis of single cell transcriptomic data from melanoma PDX demonstrated that BAHCC1 expression was the highest in “mitotic-like” melanoma cells, (Berico et al., 2021). “Mitotic” melanoma cells have been found in metastatic SKCM and UVM biopsies and are characterized by the expression of E2F-dependent genes. E2F transcription factors are known to promote melanoma progression and metastasis (Ma et al., 2008), (Alla et al., 2010). These data are consistent with the fact that our *in vitro* and *in vivo* functional studies showed that BAHCC1 was involved in melanoma cell proliferation and invasion by regulating a set of E2F/KLF-dependent genes. Although MITF^LOW^ cells have significantly reduced SE17q25 activity and BAHCC1 expression, we observed that they also depend on SE17q25 and BAHCC1, whose expression may be differently regulated by other TFs important for the mesenchymal state such as AP-1, FOSL2 or TEAD4 that bind within SE17q25 (Verfaillie et al., 2015), (Fontanals-Cirera et al., 2017), (Wouters et al., 2020) (Mauduit et al., 2021).

BAH domains are known to bind post-translation modifications (PTMs) of histones such as H3K27me3 (Zhao et al., 2016), (Fan et al., 2020) and H4K20me2 (Kuo et al., 2012), (Dai et al., 2018). A previous study characterized the BAHCC1 BAH domain showing that it bound H3K27me3 and that BAHCC1 interacted with Histone de-acetylases (HDACs) and SAP30BP proteins of the polycomb repressor complex 1 (PRC1) in Acute Myeloid Leukemia (AML) to repress a large number of genes involved in myeloid differentiation and promote cell proliferation (Fan et al., 2020). In melanoma, BAHCC1 did not show a predominant repressive role, as an equivalent number of genes were up and down regulated after BAHCC1 KD, and it was recruited to active enhancers and promoters marked by H3K27ac. These observations support the idea that BAHCC1 activity in melanoma does not depend on BAH-H3K27me3 interactions, but by alternative mechanisms. Indeed, our data demonstrated a novel interaction between BAHCC1 and the SWI/SNF chromatin remodeling complexes, occurring through the BAHCC1^BAH^ domain. We suppose that BRG1 may act upstream to remodel the chromatin, creating the nucleosome-depleted regions to be occupied by BAHCC1. Alternatively, BAHCC1 may be recruited to chromatin via interaction with transcription factors such as those of the KLF/SP/E2F families to regulate their target genes. Moreover, the N-terminal region of BAHCC1, containing the CC domain, appears to be important for the recruitment of BAHCC1 to a set of genes involved in cell proliferation and DNA repair again suggesting that the BAH domain is not the only determinant of genomic recruitment. We therefore propose that either or both of the above mechanisms drive BAHCC1 genomic recruitment in melanoma in stark contrast to the BAH-H3K27me3 pathway in AML.

Metastatic melanoma is characterized by the overexpression of genes involved in the DNA damage response (DDR), which makes this tumor stage highly refractory to chemo- and radiotherapies (Kauffmann et al., 2008). DDR is essential for maintaining genomic stability to face genotoxic stress resulting from environmental and endogenous DNA damage. In the DDR, ATM, the pivotal mediator of genotoxic stress, phosphorylates histone H2AX to γH2AX to generate docking sites for proteins involved in DNA break repair. ATM also links DNA damage to the cell cycle by controlling key DNA damage checkpoints to regulate DNA break repair directly or indirectly though the control of cell cycle checkpoints. On the other hand, PARP1 is able to bind to damaged DNA sites to promote the PARylation of surrounding proteins, thus creating new scaffolds for the recruitment of DNA repair proteins involved in an alternative DNA break repair pathway. This dual DNA repair mechanism ensures that DNA breaks are repaired efficiently. However, the loss of one DNA repair pathway results in increased reliance on the other that is not essential under normal settings. Therefore, a high response rate to the PARP inhibitor Olaparib was found in patients with metastatic prostate (Mateo et al., 2015) or gastric (Bang et al., 2015) cancers harboring low expression or mutations of ATM. Since ATM was one of the E2F-dependent genes strongly regulated by BAHCC1, we wondered if this dependence could be exploited therapeutically. Interestingly, BAHCC1 KD cooperates with PARP inhibition by Olaparib to impact cell survival, associated with increased apoptosis. In addition to their potential clinical application, these data clearly demonstrate the involvement of BAHCC1 in the control of genes involved in DDR in melanoma cells and suggest that metastatic melanoma upregulates BAHCC1 to promote genomic stability and cellular fitness necessary to sustain high mitotic rates. Finally, the presence of a gene expressed in both cutaneous and uveal melanoma cells makes it possible to envisage common treatments for these two cancers.

## Acknowledgments

We thank the IGBMC facilities, in particular Paola Rossolillo and Karim Essabri for the isolation of BAHCC1 cDNA, and Dr Catherine Birck and Dr Nathalie Troffer-Charlier for the purification of the BAH and Tudor domains. We thank Prof D. Lipsker and the staff of the Strasbourg University Hospital dermatology clinic for tumour sections and Prof. G. Ghanem and Prof. J-C Marine for providing us the MM-series melanoma cultures. This study was supported by the Institut National Du Cancer (INCa) (2017-11537), the Ligue contre le Cancer (Equipe labélisée 2022, FC and ID), the Fondation ARC, and the ANR-10-LABX-0030-INRT, a French State fund managed by the Agence Nationale de la Recherche under the frame program Investissements d’Avenir ANR-10-IDEX-0002-02. Sequencing was performed by the IGBMC GenomEast platform, a member of the “France Génomique” consortium (ANR-10-INBS-0009). P.B is supported by the Ligue contre le Cancer.

## Conflict of interest

The authors declare no competing financial interest in relation to the work described

## Author Contributions

P.B., I.D and F.C. conceived the study. P.B., I.D and F.C analyzed the data. P.B. performed most of the *in vitro* functional studies, the RNA-seq and RNAScope. P.B., and M.N. performed the RT-qPCR, and the WB. S.L., G.D., T.Y and P.B., performed bioinformatic analyses. B.V performed wet lab of CUT&Tag experiments. M.C. performed the boyden-chamber experiments. G.M. supervised the in vivo studies. M. D, E.C and C.B. provided valuable materials. F.C, I.D and P.B wrote the manuscript.

## Lead Contact

Further information and requests for resources and reagents should be directed to and will be fulfilled by the Lead Contact, Frédéric Coin.

## Materials availability

Reagent generated in this study will be made available on request, but we may require a payment and/or a completed Materials Transfer Agreement if there is potential for commercial application.

## Data availability

Next generation sequencing raw and processed data have been deposited in the SuperSeries Gene Expression Omnibus under accession number GSE205463: RNA-seq GSE201702, CUT&Tag GSE205462.

## Competing Financial Interests

Authors declare no competing financial interests.

## STAR Methods

### Cell culture

Patient-derived short-term cultures MM cells have been grown in HAM-F10 (Gibco, Invitrogen) supplemented with 10% Fetal Calf Serum (FCS) and penicillin-streptomycin. 501mel and SK-MEL-28 cells were grown in RPMI w/o HEPES (Gibco, Invitrogen) supplemented with 10% FCS and gentamycin and IGR37 and IGR39 were grown in RPMI w/o HEPES supplemented with 15% of FCS and gentamycin. Uveal melanoma cells OMM1.3 and OMM2.5 were cultured respectively in DMEM (4.5g/l glucose) supplemented with 10% FCS, penicillin-streptomycin, Sodium Pyruvate, MEM essential vitamin mixture, NEAA mixture and Hepes, and RPMI w/o HEPES supplemented with 2gr/l glucose, 10% FCS and penicillin-streptomycin. U-2 OS cells were grown in DMEM/Ham-F10 (1:1) supplemented with 10% FCS and gentamicin. HEK293T cell were grown in DMEM (1g/l glucose) supplemented with 10% FCS and penicillinstreptomycin. All cell lines were grown in 5% CO2 at 37°C. Melanocyte cell line Hermes3A was grown in 10% CO2 at 37°C in RPMI w/o HEPES supplemented with 10% FCS, penicillinstreptomycin, 200nM TPA (Sigma Aldrich), 200pM Cholera Toxin (Sigma Aldrich), 10ng/mL hSCF (Life Technologies), 10nM EDN-1 (Sigma Aldrich) and 2mM Glutamine (Invitrogen). All cell lines used were mycoplasma negative.

IGR37 shRNA doxycycline-inducible were generated by transducing cells with lentiviral vectors pLT3_shCTRL or pLT3_sh4 (shBAHCC1) and pGL4.10Luc2 and further selected with 1ug/mL of puromycin and 10ug/mL of Blasticidin.

501mel HA-BAHCC1^BAH^ were generated by transducing cells with lentiviral vectors pLenti-TET-3HA-BAHCC1^BAH^ and selected with 0.5ug/mL of puromycin.

Cells carrying a doxycycline-inducible system were treated with 1ug/mL of doxycycline for at least 24h.

GapmeRs or siRNAs were transfected in cells with Lipofectamine RNAiMAX following the manufacture instructions using an oligos final concentration of 25nM and cells were harvested 48h and/or 72h after transfection.

For CRISPRi experiments, 501mel cells were co-transfected with pX629_dCas9_KRAB_mScarlet (plasmid obtained from the IGBMC BioMol service) and pCDNA-GFP (gCTRL) or pcDNA-GFP-SE17q25 (gSE17q25) using Fugene6 following manufacture instructions. Afterward, double positive GFP+/mScarlet+ cells were sorted with a FACSaria Fusion BD Biosciences Cell sorter and RNA extraction was performed 72h post sorting.

BAHCC1^TUDOR^ and BAHCC1^BAH^ were cloned into a pcDNA-FLAG vector. pcDNA-FLAG, pcDNA-FLAG-BAHCC1^TUDOR^, pcDNA-FLAG-BAHCC1^BAH^, pLenti6-GFP and pLenti6-3HA-BAHCC1^BAH-TUDOR^ vectors were transiently transfected in HEK293T cells with Lipofectamine 2000 following the manufacture instructions.

### Identification of SEs

H3K27ac ChIP-seq data from 12 different melanoma cell lines (MM001, MM011, MM031, MM034, MM057, MM074, MM047, MM087, MM099, MM118, SK-MEL-5, 501mel) and 3 melanocytes cell lines (NHEM1, NHEM2, Foreskin) were retrieved from GEO GSE60666, GSM958157 and GSE94488 and mapped to the Homo Sapiens genome (assembly hg19) using Bowtie v1.0.0 with default parameters except for “-p 3 -m 1 –strata –best –chunkmbs 128”. Normalized BigWig files were generated using Homer makeUCSCfile v4.9.1 with the following parameter’-norm 20e6’ meaning that data were normalized to 20M reads. The genome was divided into bins of 10Kb long with Deeptools multiBamSummary v2.5.0. The number of reads for each bin was computed for each sample. The following figure was made with Deeptools plotCorrelation and shows the pairwise correlation values (Spearman) for all samples of this project. Peaks were called using MACS2 with default parameters except for “-g hs -f BAM –broad –broad-cutoff 0.1”. Peaks falling into ENCODE blacklisted regions (“An Integrated Encyclopedia of DNA Elements in the Human Genome” 2012–9AD) were removed. Peaks were annotated relative to genomic features using Homer v4.9.1 (annotations got extracted from gtf file downloaded from ensembl v75). ROSE was used to differentiate SEs from typical enhancers (detected from H3K27ac data). SEs were annotated relative to genomic features using Homer v4.9.1 (annotations got extracted from gtf file downloaded from ensembl v75). Finally, SEs were filtered according to their position relative to the one of SOX10 and MITF (ChIP-seq tracks in 501mel) (Strub et al., 2011), (Laurette et al., 2015) using the bioconductor package DiffBind v1.12.3.

### RNA FISH

For the detection of BAHCC1 and SOX10, cells and paraffin tissue sections were treated following the RNAScope™ manufacture protocol. Cells and tissue samples were counterstained with DAPI and visualized using confocal microscope Spinning disk Leica CSU W1. The sequences of the probes were not provided by the manufacture.

### Protein extraction

For the production of whole cell extracts, cells were washed once with cold PBS, rinsed with a cell scraper, pelleted and resuspended in LSDB 0.5M buffer (0.5M KCl, 50mM Tris HCl pH 7.9, 20% Glycerol, 1% NP40, 1mM DTT, PIC). Afterwards cells were fully disrupted with 3 cycles of heat shock in liquid nitrogen and 37°C water bath and centrifugated 15min at 14,000rpm to pellet cell debris.

To obtain cytoplasmic and nuclear protein fractions, cells were first lysed in hypotonic buffer (10 mM Tris–HCl at pH 7.65, 1.5 mM MgCl_2_, 10 mM KCl) and disrupted by Dounce homogenizer. The cytosolic fraction was separated from the pellet by centrifugation at 4°C. The nuclear soluble fraction was obtained by incubation of the pellet in high salt buffer (final NaCl concentration of 300 mM) and then separated by centrifugation at 4°C. To obtain the nuclear insoluble fraction (chromatin fraction), the remaining pellet was digested with micrococcal nuclease and sonicated.

### Co-immunoprecipitation

Whole cell extract was prepared by resuspending cells in LSDB buffer (50mM Tris HCl pH 7.9, 20% Glycerol, 1% NP40, 1mM DTT, PIC) containing 150mM KCl, followed by sonication in Q800R3 sonicator. Between 250ug-1mg of protein extract was used to performed IP with 1-10ug of primary antibody overnight at 4°C on rotation. Following, 50ul of Dynabeads protein A/G were added to the samples for 2h at 4°C. In alternative, FLAG-tagged proteins were immunoprecipitated directly using Affinity Gel FLAG M2 conjugated beads. Beads were washed five times with TGEN buffer (20mM Tris-HCl pH 7.65, 3mM MgCl_2_, 0.1mM EDTA, 10% glycerol, 0.01% NP-40, PIC) containing 150mM NaCl. Samples were loaded on NuPage gel to perform western blot

### RNA extraction and RT-qPCR

Total RNA isolation was performed according to the manufacture protocol with NucleoSpin RNA Plus kit. RNA was retrotranscribed with Reverse Transcriptase Superscript IV and qPCR was performed with SYBR Green and monitored by LightCycler 480. Gene expression results were normalized according to four housekeeping genes (HMBS, TBP, UBC and RPL13a). Primers for RT-qPCR and ChIP-qPCR were designed using Primer-BLAST.

### Immunofluorescence

Human tissue sections were deparaffinized and dehydrated with Histosol and dilutions of ethanol 100%, 90%, 70% and 30% and rehydrated with demineralized water. Subsequently sections were boiled in Sodium Citrate buffer (0.1M Citric acid, 0.1M Sodium citrate) for 15min to unmask antigens. In parallel, 2D culture cells were grown on LAB-TEK II chamber slides and fixed with 4% formaldehyde or 100% methanol. Afterward, both tissues and cells were permeabilized and saturated in blocking buffer (1% BSA, PBS, 0.3% TritonX-100). Primary antibodies were diluted in blocking buffer and incubate ON at 4°C in wet chamber. Secondary antibody staining was carried out in blocking buffer for 1h and 30min at room temperature. Nuclei were marked with DAPI and slides were mounted with ProLong™ Gold antifade reagent before microscope image acquisition.

For γH2AX foci quantification, transfected or naïve cells were plated in 96 well plates OptiPlates-96 and eventually treated with NCS (150nM) for 1h at 37°C followed by 1h, 8h or 24h of recovery in complete medium. Afterward, cells were labeled for γH2AX following a classic immunofluorescence protocol. Image acquisition was done using high-throughput imaging system CX7 using 20X objective (50 fields per well). Image segmentation was done with HCS studio. Nuclei were identified using DAPI staining and γH2AX foci were identified within nuclei mask. Foci number and intensity were quantified automatically.

### Flow cytometry

To measure cell proliferation, cell death and senescence, cells were incubated first with 1uM of CellTrace Violet according to the manufacture instruction. At the end of transfection and/or drug treatment, cells were rinse and incubate 15min with AnnexinV-APC. Cells proliferation and cell death were detected on a FORTESSA BD Biosciences Cytofluorometer. Data were analyzed with FlowJo software.

### Cell density assay

Following transfection, between 5.10^4^ to 1.10^5^ cells were grown in 6 wells plate for up to 1 week. Afterward cells were fixed for 10min with 4% Formaldehyde solution, washed once with PBS and stained with Crystal Violet solution 0.2% for 15min at room temperature. The wells were finally washed twice with deionized water, air dried, scanned and analyzed with Fiji considering the area occupancy of the cells.

### Boyden-Chamber invasion assay

Between 1 and 2.10^5^ cells were seeded inside a Boyden Chamber insert covered with Serum free media and 4% Matrigel. The inserts were placed in 24 wells plate filled with complete medium. After 12-24h, the inserts were fixed with 4% Formaldehyde solution for 10min, gently cleaned inside with a cotton stick and stained for 15min at room temperature with a Crystal Violet solution 0.2%. Afterward the inserts were washed twice in deionized water, air dried and photos were collected using an EVOS xl Core microscope. The pictures were analyzed with Fiji considering the area occupancy of the cells.

### CUT&Tag and deep sequencing

Assays were performed following the manufacturer’s instructions. Briefly, 5.10^5^ cells per condition were used. The cells were washed 2 time before the binding on Concavalin A beads and then incubated overnight with BAHCC1 (BAHCC1^WT^) or HA (BAHCC1 ^TUDOR-BAH^) primary antibodies at the recommended dilution (1:50) or without antibody (negative control). The next day the corresponding secondary antibody, a guinea pig Anti-rabbit antibody was used following a 1:100 dilution in digitonin buffer and incubated at room temperature for 1h. The CUT&Tag-IT™ Assembled pA-Tn5 Transposomes were incubated for 1h at room temperature before tagmentation. Cells were resuspended in Tagmentation buffer and incubated at 37°C for 1h, then the tagmentation process was stopped by addition of EDTA and SDS. Protein digestion was performed by the addition of 80ug/mL of proteinase K and incubated at 55°C for 60min. DNA was retrieved on DNA purification columns provided by the manufacturer. Library preparation and PCR amplification were done using the Kit primers and purified by 2 successive washes with SPRI beads. Samples were subjected to paired-end sequencing by the IGBMC GenomEast platform on Illumina HiSeq 4000 instrument.

### Deep sequencing analysis

ChIP-seq data for BRG1 (Laurette et al., 2015), H3K27ac and ATAC-seq (Fontanals-Cirera et al., 2017) and CUT&Tag-seq for BAHCC1 (this work) were analyzed as previously described (Berico et al., 2021). Briefly, after reads mapping onto hg19 human genome, peak calling was performed using MACS2 according to specific negative control inputs (Zhang et al., 2008). Peak annotation was carried out using HOMER and ChIPseeker. Peak genome distribution and correlation between BAHCC1 and BRG1, H3K27ac and ATAC was performed using deepTool2. Peak enrichment for BRG1, H3K27ac, ATAC and BAHCC1 was done using seqMINER. Top 500 macs peaks summits for BAHCC1 alone or co-bound by BAHCC1 and BRG1 (identified with seqMINER) were subsequently extended of 100bp upstream and downstream using “bedtools slop” followed by extraction of FASTA format sequences with “bedtools getfasta”. DNA binding motif analysis was carried out with Simple Enrichment Analysis (SEA) (Bailey and Grant, 2021) and by the pipeline “peak-motifs” available online as part of the Regulatory Sequence Analysis Tools (RSAT) web server using +/− 400bp around the center to cut peak sequences and the non-redundant vertebrate Jaspar core database for motif comparison.

### Bulk RNA-seq analysis

Reads were preprocessed to remove adapter and low-quality sequences (Phred quality score below 20). After this preprocessing, reads shorter than 40 bases were discarded for further analysis. These preprocessing steps were performed using cutadapt version 1.10. Reads were mapped to rRNA sequences using bowtie version 2.2.8 and reads mapping to rRNA sequences were removed for further analysis. Reads were mapped onto the hg19 assembly of Homo sapiens genome using STAR version 2.5.3a. Gene expression quantification was performed from uniquely aligned reads using htseq-count version 0.6.1p1, with annotations from Ensembl version 75 and “union” mode. Only non-ambiguously assigned reads have been retained for further analyses. Read counts have been normalized across samples with the median-of-ratios method proposed by Anders and Huber (Anders and Huber, 2010) to make these counts comparable between samples. Comparisons of interest were performed using the Wald test for differential expression proposed by Love (Love et al., 2014) and implemented in the Bioconductor package DESeq2 version 1.16.1. Genes with high Cook’s distance were filtered out and independent filtering based on the mean of normalized counts was performed. P-values were adjusted for multiple testing using the Benjamini and Hochberg method (Benjamini and Hochberg, 1995). iRegulon plugin of Cytoscape was used to analyze the co-deregulated genes between iBAHCC1 and iBRG1.

### Single cell data analysis

Expression matrix with row reads counts for the single cell experiment was retrieved from GEO GSE116237, GSE115978, GSE151091 and GSE138665 datasets. Then, data were normalized and clustered using the Seurat software package version 3.1.4 in R version 3.6.1. Data were filtered and only genes detected in at least 3 cells and cells with at least 350 detected genes were kept for further analysis. Expression of 26,661 transcripts in single cells was quantified. To cluster cells, read counts were normalized using the method “LogNormalize” of the Seurat function NormalizeData. It divides gene expression counts by the total expression, multiplies this by a scale factor (10,000 was used), and log-transforms the result. Then, 2000 variable features were selected with the variance stabilizing transformation method using the Seurat function FindVariableGenes with default parameters. Integrated expression matrices were scaled (linear transformation) followed by principal component analysis (PCA) for linear dimensional reduction. The first 20 principal components (PCs) were used to cluster the cells with a resolution of 0.5 and as input to tSNE to visualize the dataset in two dimensions. The Bioconductor package AUCell v 1.6.1 was used to assess whether some cells from the different datasets were enriched in gene sets of interest.

### Other publicly available datasets used in this work

BAHCC1 RNA expression was quantified in available datasets including GEO GSE12391, GSE80829, GSE98394, GSE114445, GSE46517, GSE8401 and in the CCLE portal. TCGA and GTEx data were obtained through UCSC Xena browser, cBioPortal and Gene Expression Profiling Interactive Analysis 2 (GEPIA2). PCA profile of TCGA and melanoma cell lines according to their phenotypic profile was obtained from the Graeber Lab software

### Xenograft studies

Animal experiments were approved by the French « Ministère de l’enseignement supérieur, de la recherche et de l’innovation » with the APAFiS n° C6721840. Immunodeficient mice were purchased from Charles River Laboratory and maintained in the animal facility germ-free of the PHENOMIN-ICS. 3.10^6^ IGR37 cells were resuspended in 100ul of PBS + 100ul of Culturex^®^ Basement Membrane Extract and injected in the flank of the animals. In parallel, mice were feed with food supplied with 200mg/kg doxycycline to induce the shRNAs expression in the injected cells. Tumor progression was quantified once per week through luciferase radiance bioluminescence with IVIS imager until the primary tumor size reached a size of 1cm^3^. Briefly, mice were injected with 100ul of D-Luciferin (150mg/kg in PBS) intraperitoneal and anesthetized with 4% isoflurane before radiance measurement. Tumor volume (*V*) was measured every 3 days using the formula 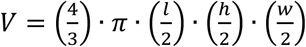 were *l* is length, *h* is height and *w* is width of the tumor.

Mice tumors were dissected and dissociated in single cell suspension using gentleMACS™ Dissociator in combination with gentleMACS C-tubes and Human Tumor dissociation kit following manufacture instructions. Afterward, single cell suspension tumors were processed for protein and RNA extraction as described above

### Statistical analysis

Statistical analysis was mostly performed using Prism 9. Briefly, for absolute quantification comparison, Student’s t-test and ordinary one-way ANOVA unpaired were used; paired test was used for relative quantification comparisons. Grouped sample analysis was carried out through two-way ANOVA test. Kaplan-Meier survival curves were analyzed using Mantel-Cox test. For correlation analysis, Spearman analysis was performed together with linear regression curve fit. Statistical analysis on RNA-seq and ChIP-seq is listed in their dedicated sections. For Venn diagram statistic, hypergeometric test was performed using Nemates software (nemates.org). P values are represented as ns (p>0.05), * (p<0.05), ** (p<0.01), *** (p<0.005) and **** (p<0.001).

---

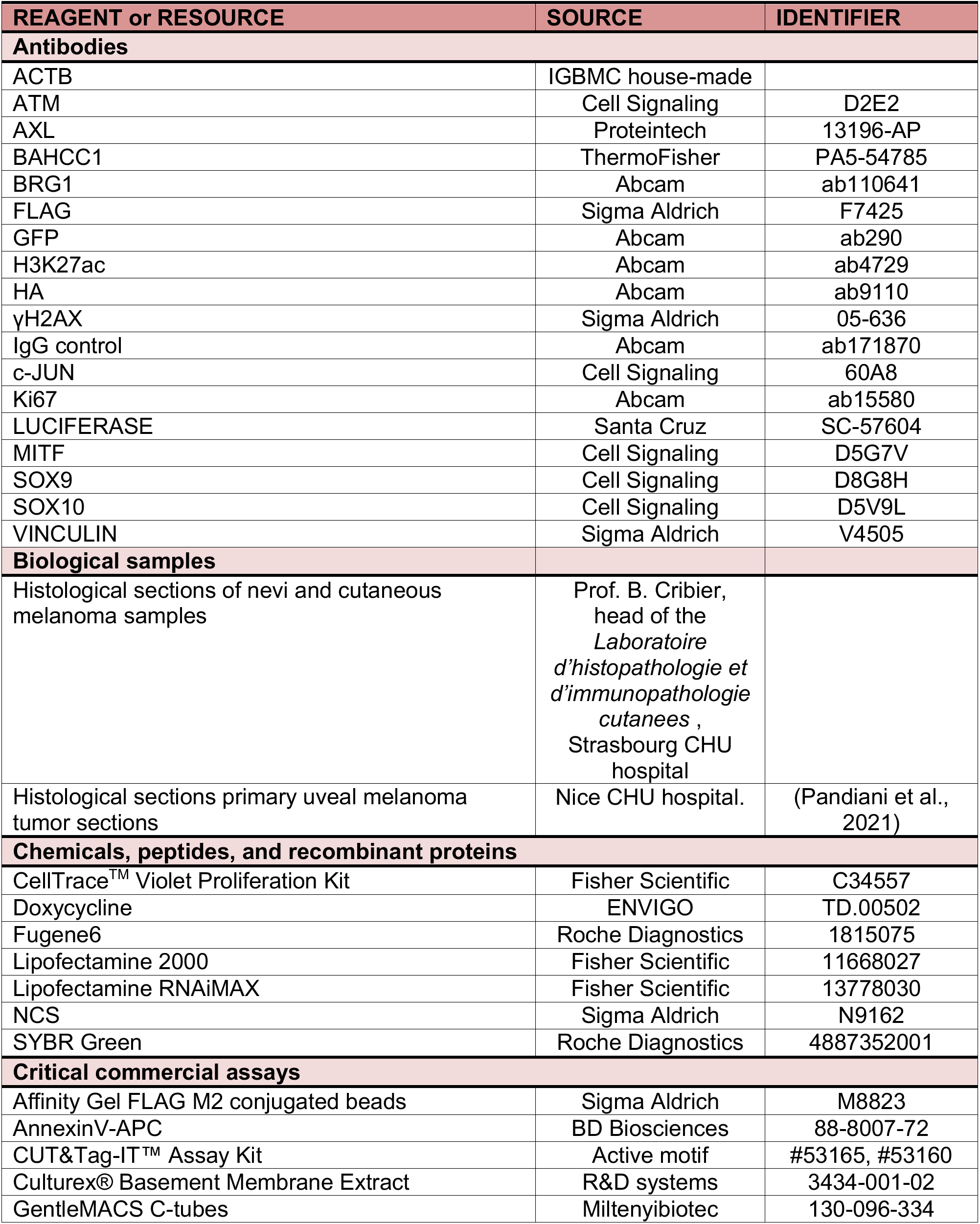
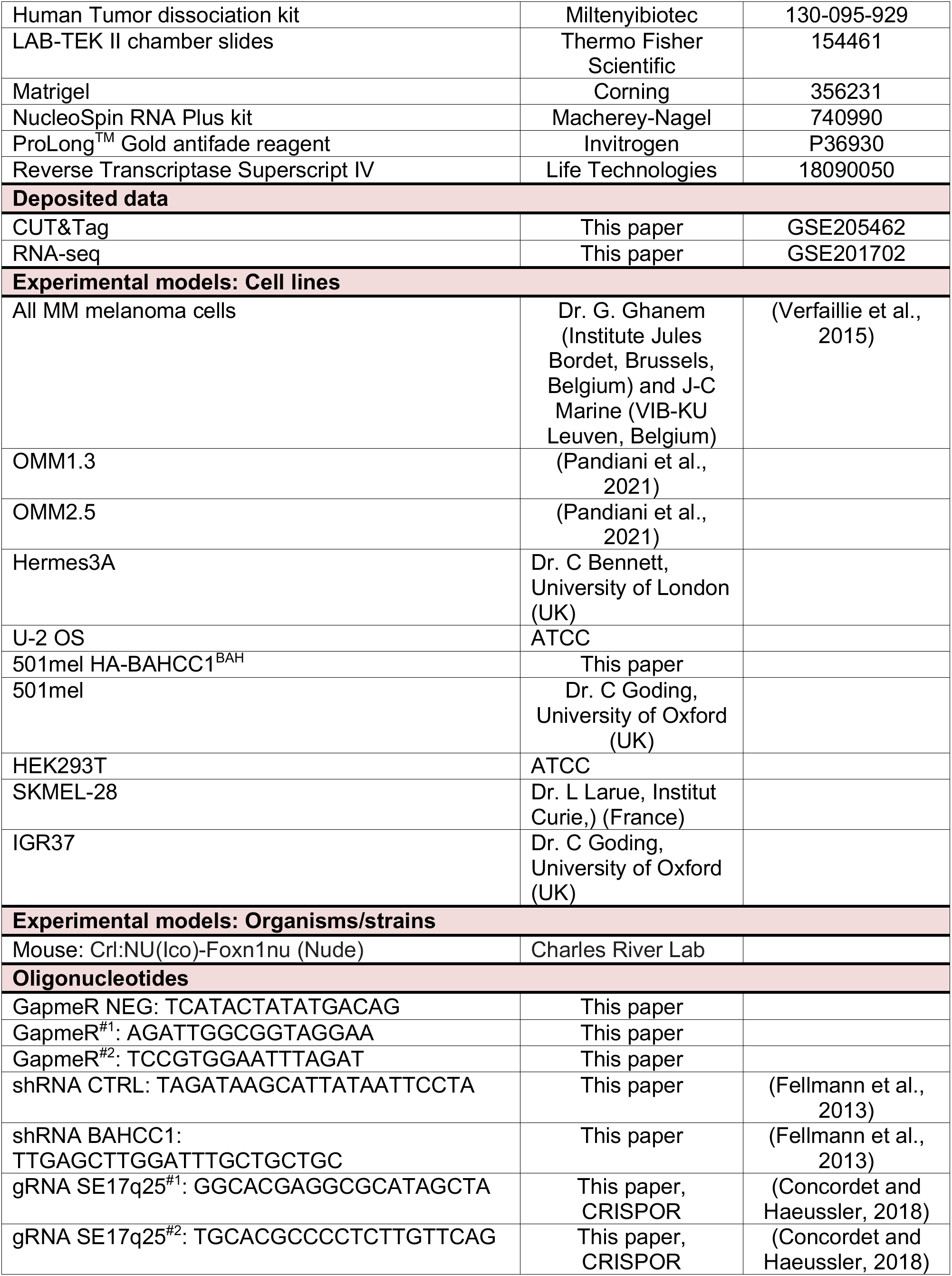
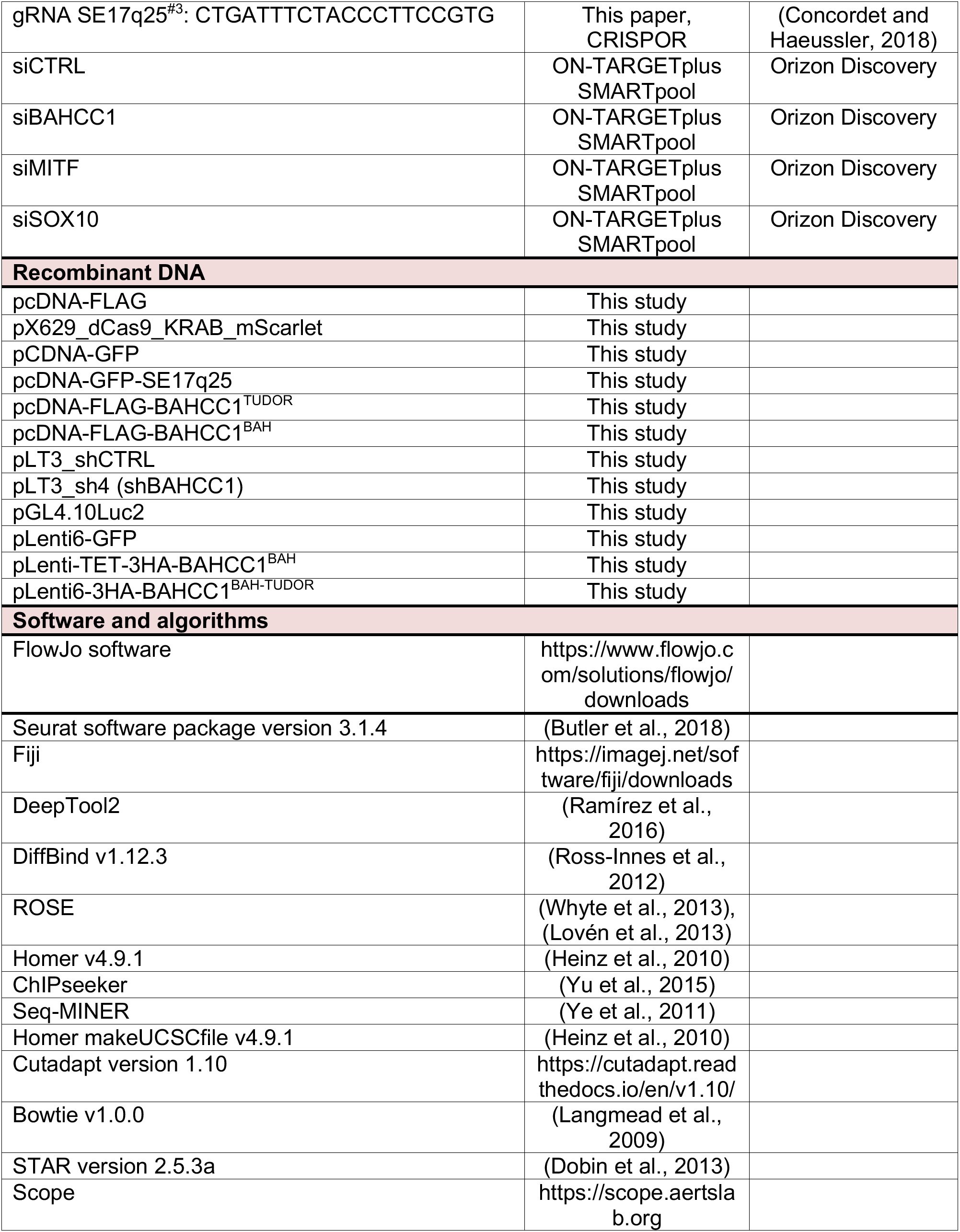
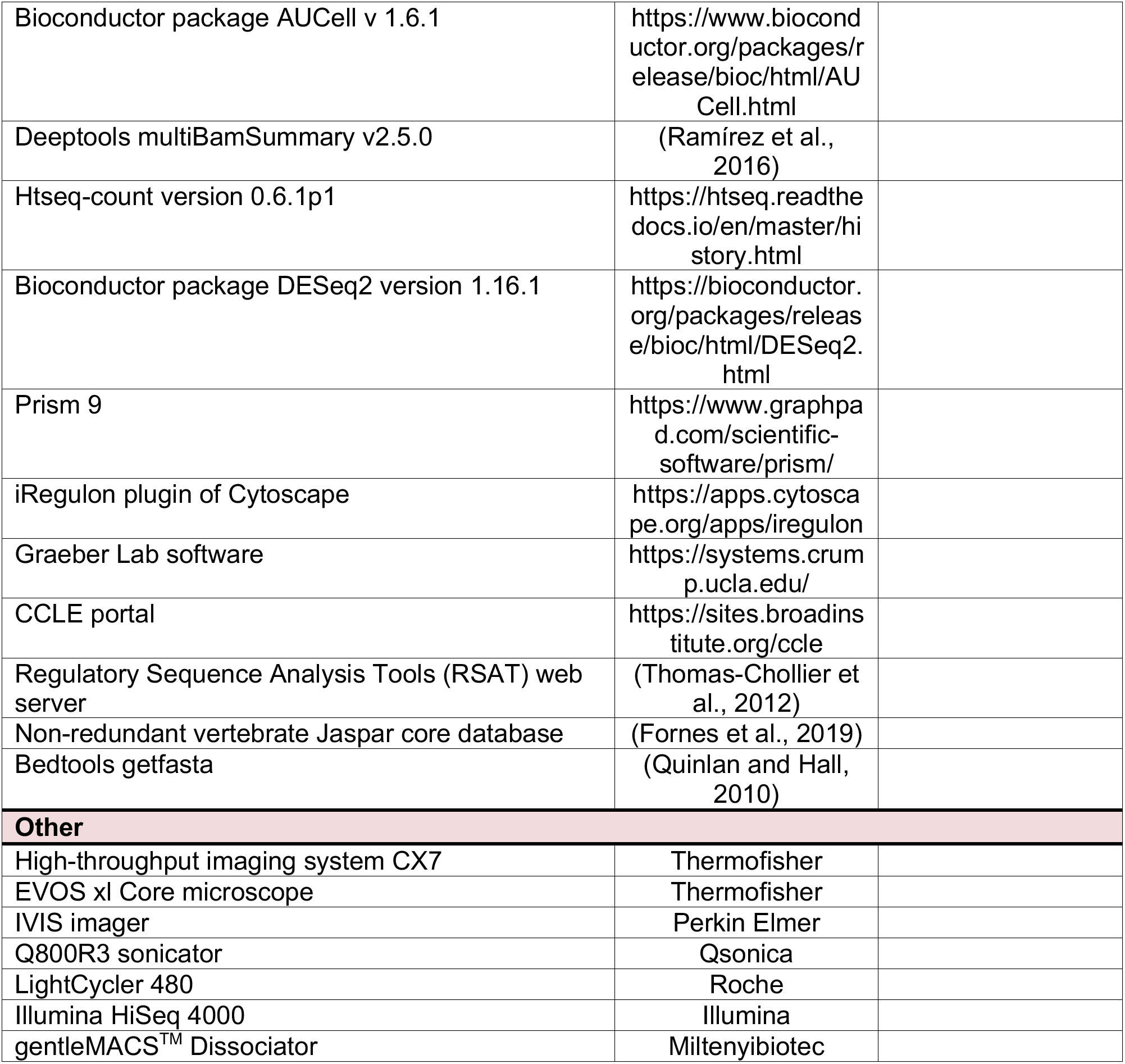

**Ext. Figure 1:**
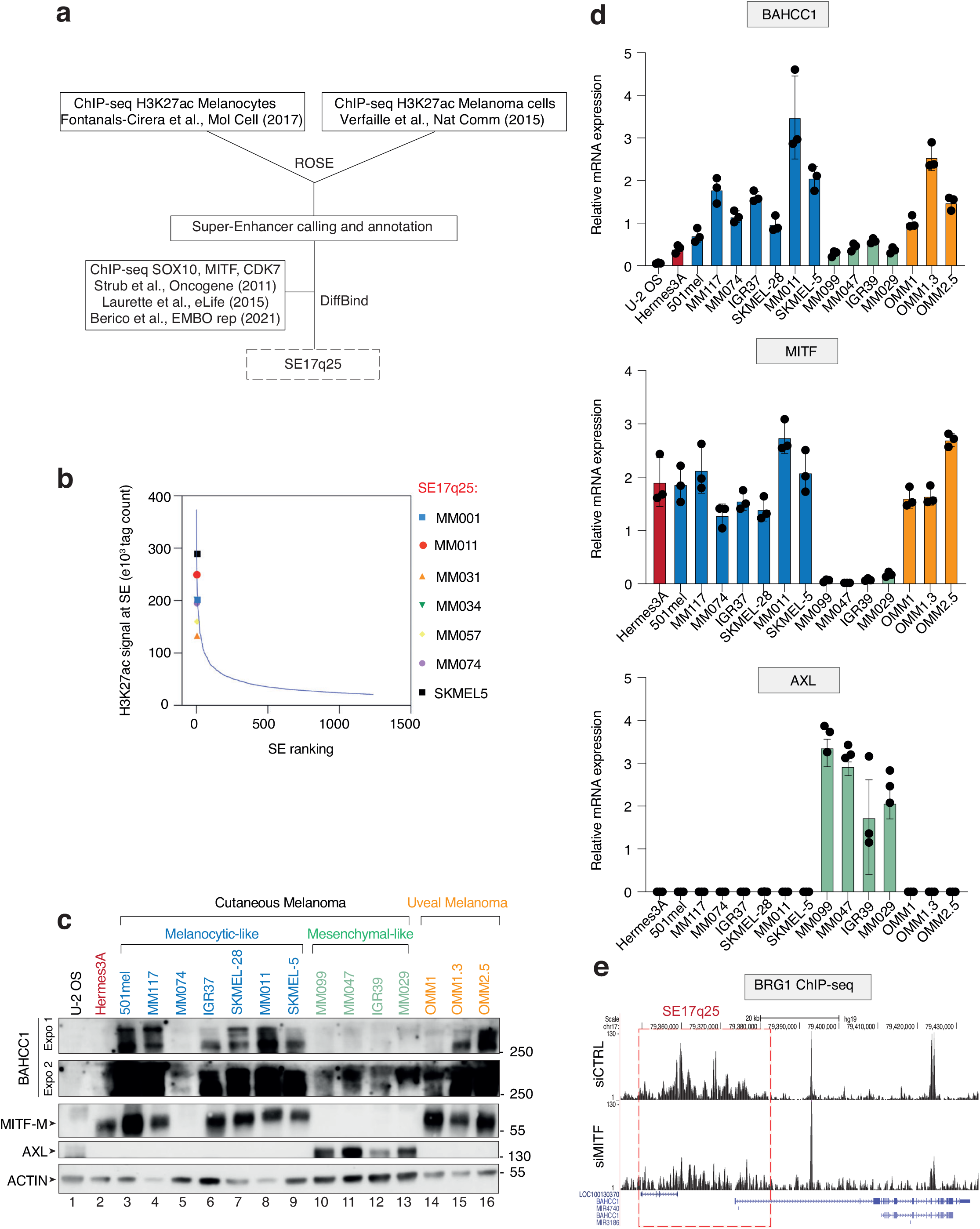
BAHCC1 overexpression in melanoma cells. **a**. Table summarizing the different cutaneous melanoma cells used in this study. The driver mutation and the cellular phenotype are indicated. MM cells are patient-derived short-term melanoma cultures. **b.** Schematic representation of the SE pipeline analysis used to identify SE17q25. **c.** Dot plot representing the H3K27ac signal at SEs as a function of the ROSE ranking. We used the distribution of SEs in MM011 cells as a representative distribution. In different shape and colors are the specific positions of SE17q25 in the corresponding melanoma cell lines. **d**. Immunoblot against BAHCC1, MITF, AXL and ACTIN in U-2 OS, immortalized melanocyte Hermes3A and several SKCM and UVM cell lines as indicated. **e**. Relative expression of BAHCC1, MITF and AXL determined by RT-qPCR in a large panel of melanoma and non-melanoma cell lines as indicated. Bars represent mean values of three different experiments (Biological triplicates) (+/− SEM). **f**. BRG1 ChIP-seq tracks in 501mel treated with siCTRL or siMITF, scaled in the region of SE17q25 (red box). The decommission of SE17q25 is illustrated by the loss of BRG1 binding. Values were retrieved from the GEO dataset GSE61967.

**Ext. Figure 2:**
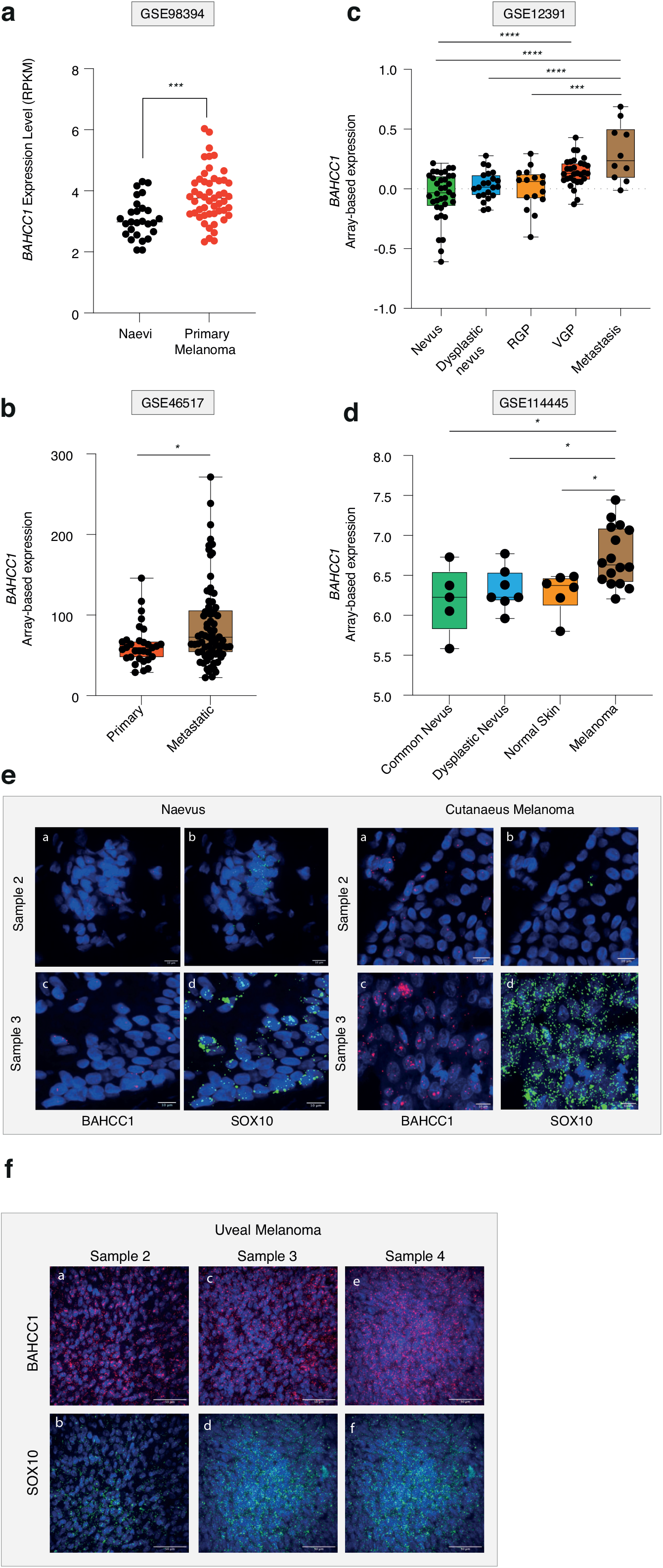
BAHCC1 expression increases during melanoma progression. **a**. BAHCC1 expression in human nevi (n=27) and primary melanomas (n=51). Values were retrieved from the GEO dataset GSE98394. **b**. BAHCC1 expression in primary (n=31) and metastatic (n=73) melanoma. Values were retrieved from the GEO dataset GSE46517. **c**. Expression levels of BAHCC1 in human biopsies of nevus (n=36), dysplastic nevus (n=22), melanoma radial growth phase (RGP) (n=16), vertical growth phase (VGP) (n=30) and metastatic melanoma (n=10). Values were retrieved from the GEO dataset GSE12391. **d**. BAHCC1 expression in normal skin (n=6), common nevus (n=5), dysplastic nevus (n=7) and melanoma (n=16). Values were retrieved from the GEO dataset GSE114445. **e-f**. RNA FISH for SOX10 and BAHCC1 in human tissue samples of nevi (n=2), cutaneous (n=2) and uveal melanomas (n=3, same samples than (1)). Scale bar is indicated for each image.

**Ext. Figure 3:**
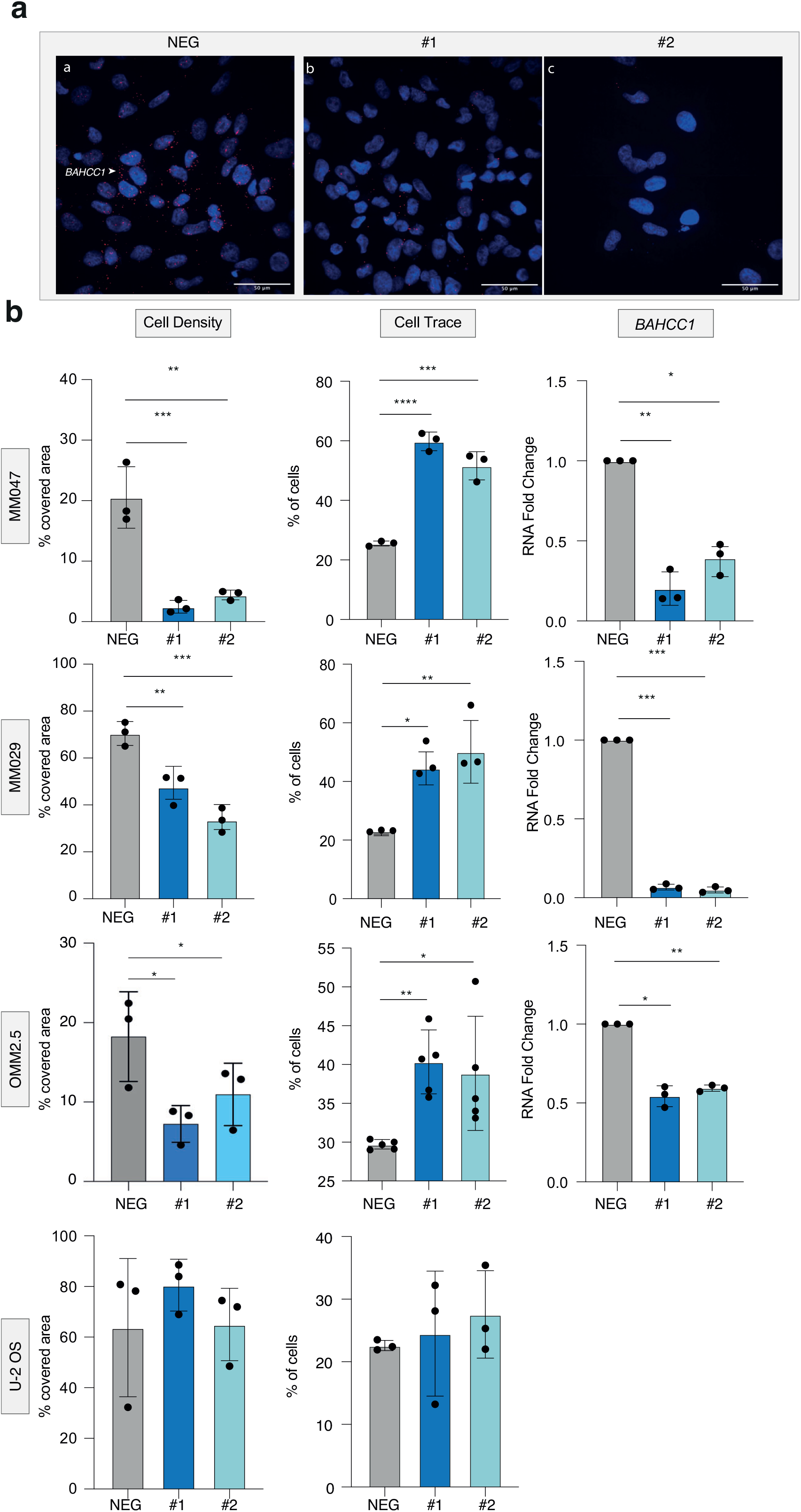
BAHCC1 depletion impairs SKCM and UVM cells proliferation. **a**. RNA FISH was performed for BAHCC1 in 501mel and MM047 transfected with GapmeR^NEG^ (NEG), GapmeR^#1^ (#1) and GapmeR^#2^ (#2). Scale bars are indicated. **b**. **Left panel**; Cell coverage quantification of crystal violet staining of cells transfected with GapmeR^NEG^ (NEG), GapmeR^#1^ (#1) and GapmeR^#2^ (#2) in SKCM cells (MM047 and MM029) or UVM cells (OMM2.5). **Middle panel;** CellTrace staining was measured by FACS in the cells used in the left panel and results are represented as % of slow proliferative cells considering an arbitrary threshold between 20-30% in the GapmeR^NEG^ (NEG) control. **Right panel;** Relative BAHCC1 expressions upon transfection with GapmeR^NEG^ (NEG), GapmeR^#1^ (#1) and GapmeR^#2^ (#2) were measured by RT-qPCR in the cells used in the left panel (with the exception of U-2 OS cells that do not express BAHCC1). Bars represent mean values of three different experiments (Biological triplicates) (+/− SEM). Two-way ANOVA using Šídák’s multiple comparisons test.

**Ext. Figure 4:**
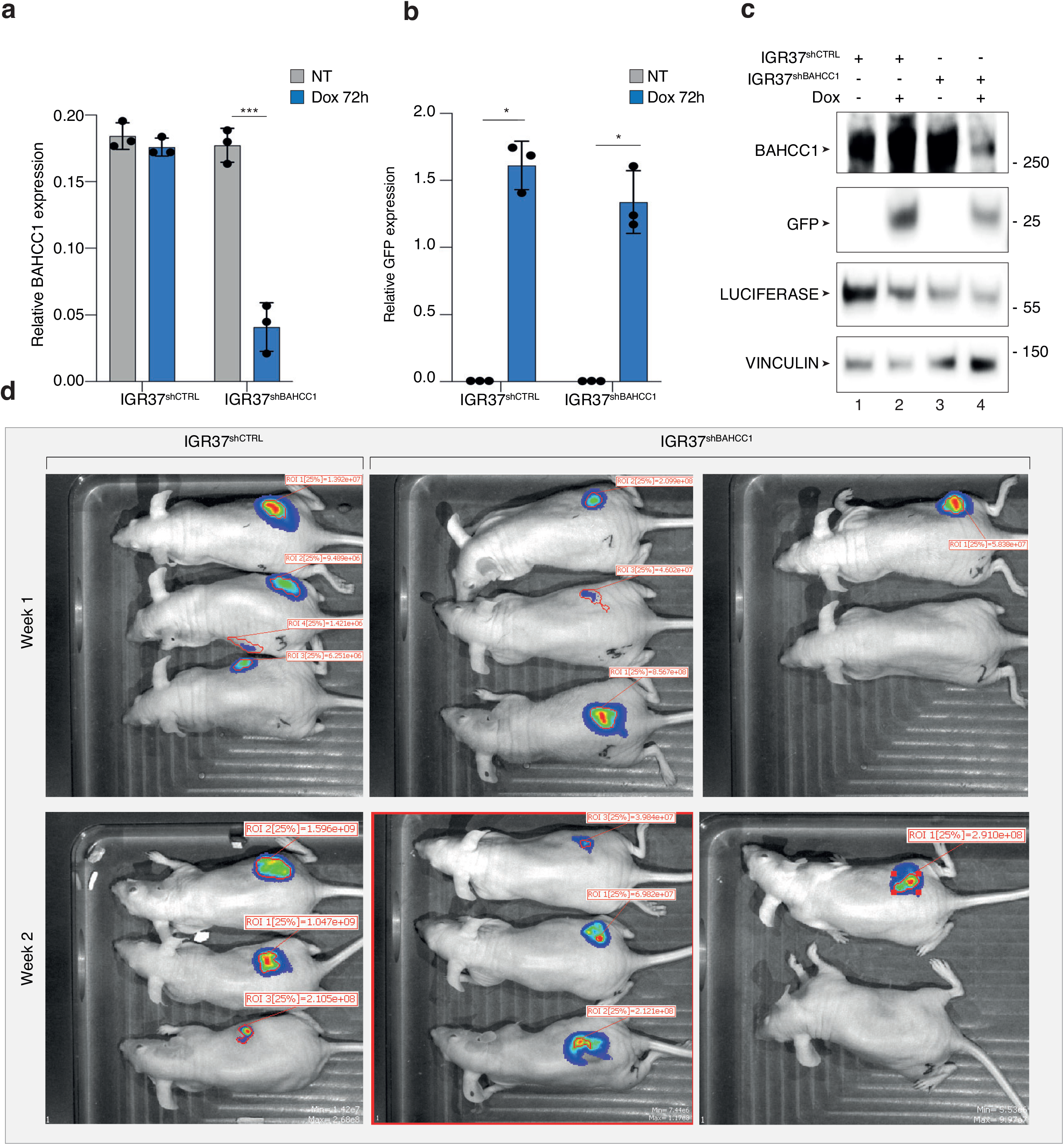
CDXs mouse models of BAHCC1 depletion. **a-b**. BAHCC1 **(a)** and GFP **(b)** relative expression was determined by RT-qPCR in IGR37^shCTRL^ and IGR37^shBAHCC1^ melanoma cells non treated (NT) or treated with doxycycline (Dox) for 72h *in vitro*. Bars represent mean values of three different experiments (Biological triplicates) (+/− SEM); Two-way ANOVA using Šídák’s multiple comparisons test. **c**. Total extracts from IGR37^shCTRL^ and IGR37^shBAHCC1^ cells treated or not with Dox were resolved by SDS-PAGE and proteins were immunoblotted as indicated. Luciferase is expressed in all conditions for *in vivo* IVIS imaging. **d**. IVIS photos of mice bearing IGR37^shCTRL^ (n=3) or IGR37^shBAHCC1^ (n=4) derived tumors. Radiance colorimetric scale is normalized between the different samples.

**Ext. Figure 5: BAHCC1 localizes at the promoter of E2F/KLF dependent genes**

**a**. Immunostaining of BAHCC1 in 501mel cells using an anti-BAHCC1 antibody together with DAPI staining. Scale bars are indicated.

**b**. Immunoblot of the indicated proteins in 501mel protein sub fractions corresponding to cytoplasm (Cyto), nuclear soluble (NS) and nuclear insoluble (NI). BRG1 and Actin were used as positive controls for NS/NI and Cyto respectively.

**c**. GO analysis of the 200 co-downregulated (Top) and 82 co-upregulated (Bottom) genes between 501mel and MM047 upon BAHCC1 KD. The first 10 most significant annotation groups are listed from top to bottom according to the -log_10_(q-value).

**d**. Pie chart displaying the distribution of 31,280 BAHCC1 peaks identified in this work with respect of genomic annotations.

**e**. BAHCC1 expression among the different transcriptional states previously identified in Rambow melanoma PDXs (n=674 cells) (2), (3).

**f**. Heatmap representing the regulon activity of E2F, Sp/KLF transcription factor family members obtained by SCENIC analysis in the different transcriptional states of Rambow PDX described in **(e)**.

**g**. BAHCC1 single cell expression retrieved from the GEO dataset GSE138433 and GSE72056 was separated in two groups of “Slow” and “Fast” cell cycle cells based on cell enrichment of 95 cell cycle genes identified by Tirosh and colleagues (4) (Ext. Table 6).

**h.** SKCM (n=469) and UVM (n=81) Spearman correlation between BAHCC1 expression and Tirosh cell cycle signature (95 genes). Values were retrieved from TCGA. Analysis was done using GEPIA2.

**Ext. Figure 6:**
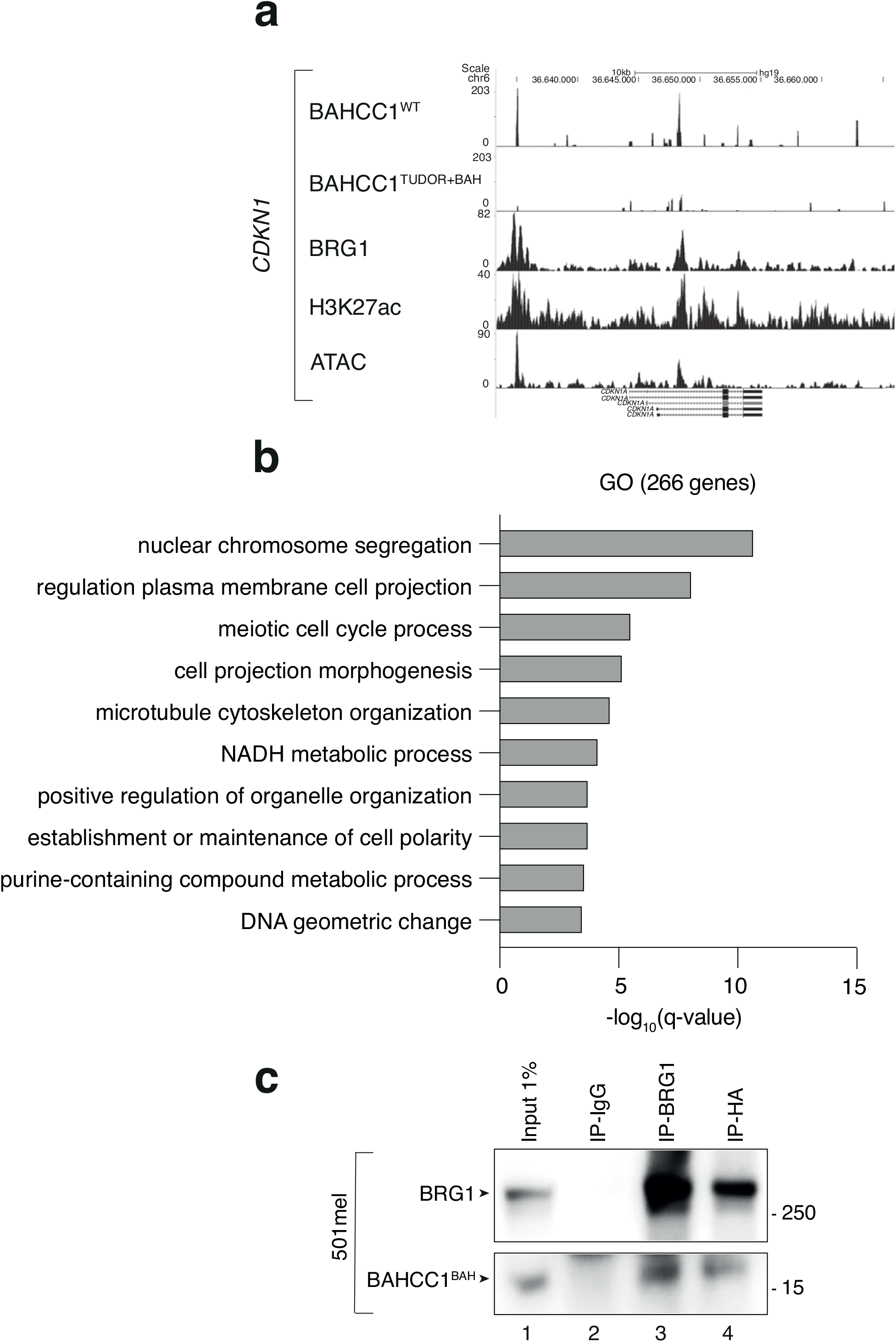
BAHCC1 and BRG1 co-regulate expression of cell cycle and DNA repair genes. **a**. Captures of the UCSC genome browser (GRCh38/hg19) showing the ChIP-seq profiles of BAHCC1^WT^, BAHCC1 ^TUDOR-BAH^, BRG1, H3K27ac and ATAC-seq in 501mel corresponding to the promoter regions of CDKN1. RefSeq annotated genes are displayed the bottom **b**. Gene annotation analysis of the 266 genes co-regulated by BRG1 and BAHCC1 in 501mel. The histogram shows the top 10 deregulated biological pathways according to the -log_10_(q-value). **c**. Following the stable overexpression of a HA-BAHCC1^BAH^ domain in 501mel cells, immunoprecipitation was carried out using anti-IgG (control), anti-BRG1 and anti-HA antibodies. Following SDS-PAGE, proteins were immunoblotted as indicated. Molecular sizes are indicated.

**Ext. Figure 7:**
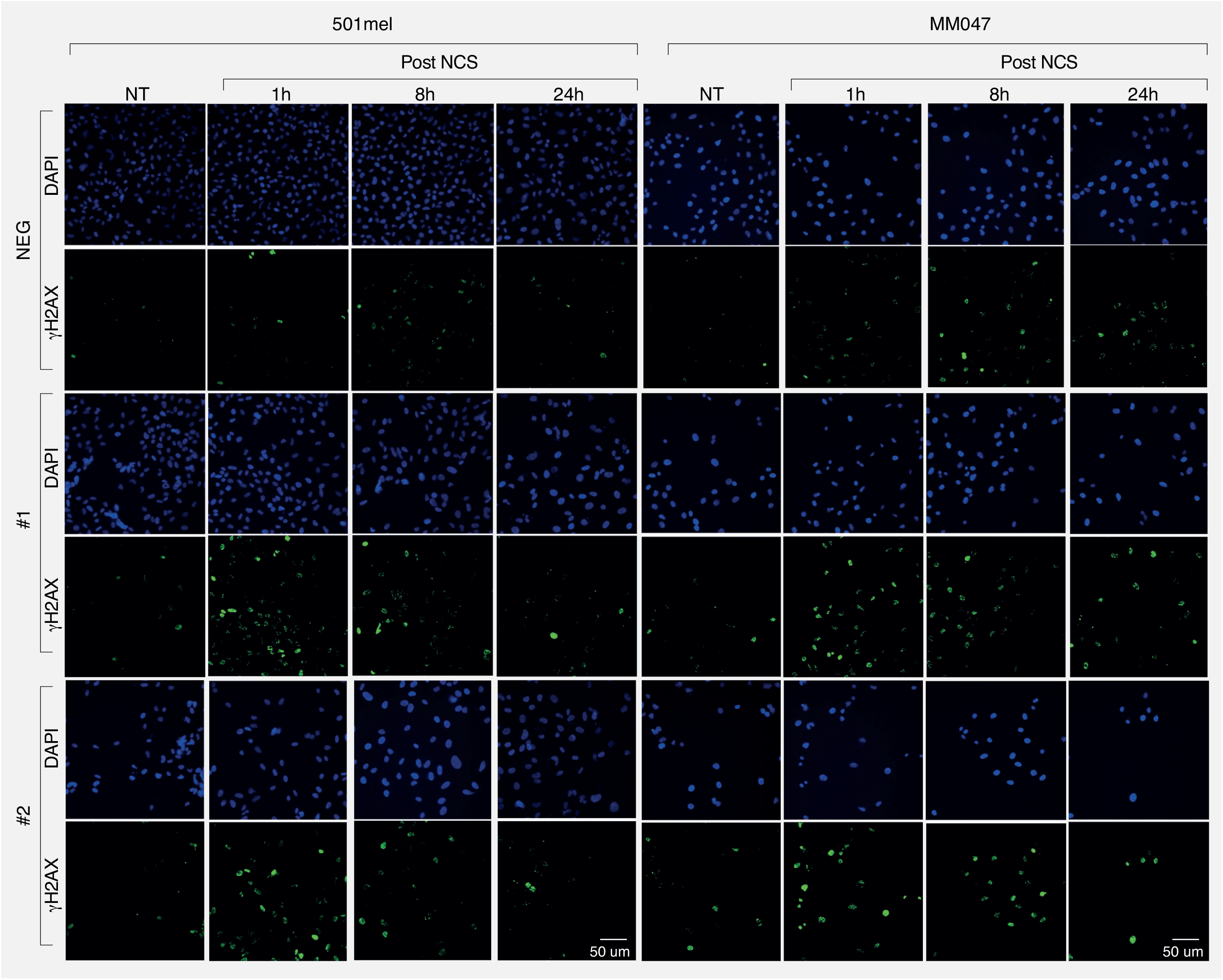
BAHCC1 depletion impairs double-stranded DNA repair. Immunofluorescence for γH2AX in 501mel and MM047 in basal conditions or upon DNA damage in presence or absence of BAHCC1. Images are representative of the quantification of Figure 7. DAPI staining is shown. Scale bars are shown.

**Extended Table 1: Melanoma cells used in this study**

The melanoma cells, their driver mutations and phenotypes are indicated.

**Extended Table 2: Putative SEs identified in melanoma cells**

The top 19 putative SEs identified in 10 different melanoma cells, including the 501mel melanoma cell line and MM cells, are shown. SE17q25 is highlighted in red and is among the top 19 SEs of the 10 cells considered.

**Extended Table 3: Down-regulated genes**

A list of genes down-regulated both in 501mel and MM047 after BAHCC1 KD is provided page 1. A list of down-regulated genes both in 501mel after BRG1 KD and after BAHCC1 KD in 501mel and MM047 is provided page 2.

**Extended Table 4: BAHCC1 binding sites**

RSAT was carried out on the BAHCC1 occupied sites according to MACS2 ranking on the top peaks (page 1), on BAHCC1 N-ter dependent peaks (page 2), on TSS of up-regulated genes in BAHCC1^KD^ cells (page 3) and on TSS of down-regulated genes on BAHCC1^KD^ (page 4).

**Extended Table 5: Gene cell cycle**

List of 95 melanoma cell cycle genes identified by Tirosh and colleagues and used in this work to separate in “Slow” or “Fast” cell cycle either single cells or bulk RNA-seq data.

**Extended Table 6: BAHCC1^WT^ *vs*. BAHCC1 ^TUDOR-BAH^ binding sites**

A list of genes bound by BAHCC1^WT^ but not by BAHCC1^TUDOR-BAH^ as determined by CUT&Tag is provided page 1. A list of down- and up-regulated genes belonging to this cluster is provided page 2.

## Notes

### Competing Interest Statement

The authors have declared no competing interest.

